# A Peptide Triple Agonist of GLP-1, Neuropeptide Y1, and Neuropeptide Y2 Receptors Promotes Glycemic Control and Weight Loss

**DOI:** 10.1101/2022.11.07.515458

**Authors:** Kylie S. Chichura, Clinton T. Elfers, Therese Salameh, Varun Kamat, Oleg G. Chepurny, Aelish McGivney, Brandon T. Milliken, George G. Holz, Sarah V. Applebey, Matthew R. Hayes, Ian R. Sweet, Christian L. Roth, Robert P. Doyle

**Affiliations:** Syracuse University, Department of Chemistry, 111 College Place, Syracuse, NY 13244, USA; Seattle Children’s Research Institute, 1900 Ninth Ave, Seattle WA 98101, USA; University of Washington, Diabetes Research Institute and Division of Metabolism, Endocrinology, and Nutrition, Seattle, WA 98195, USA; State University of New York, Upstate Medical University, Department of Medicine, Syracuse, NY 13210, USA; State University of New York, Upstate Medical University, Department of Pharmacology, Syracuse, NY 13210, USA; Department of Psychiatry, Perelman School of Medicine, University of Pennsylvania, Philadelphia, PA 19104, USA; Seattle Children’s Hospital, University of Washington, Department of Pediatrics, Seattle, WA 98105, USA

**Keywords:** GLP-1R, Y1-R, Y2-R, biased agonism, T2DM, obesity, insulin secretion, metabolism

## Abstract

Mechanisms underlying long-term sustained weight loss and glycemic normalization after obesity surgery include changes in gut hormone levels, including glucagon-like peptide 1 (GLP-1) and peptide YY (PYY). We demonstrate that two peptide biased agonists (GEP44 and GEP12) of the GLP-1, neuropeptide Y1, and neuropeptide Y2 receptors (GLP-1R, Y1-R, and Y2-R, respectively) elicit Y1-R antagonist-controlled, GLP-1R-dependent stimulation of insulin secretion in both rat and human pancreatic islets, thus revealing the counteracting effects of Y1-R and GLP-1R agonism. These agonists also promote insulin-independent Y1-R-mediated glucose uptake in muscle tissue *ex vivo* and more profound reductions in food intake and body weight than liraglutide when administered to diet-induced obese rats. Our findings support a role for Y1-R signaling in glucoregulation and highlight the therapeutic potential of simultaneous receptor targeting to achieve long-term benefits for millions of patients.

## INTRODUCTION

The pathophysiology of obesity is driven by the dysregulation of numerous interrelated pathways. Thus, interventions that are effective at treating obesity will most likely be those that target multiple receptors in complementary neurocircuits and regulate energy balance. Patients who have undergone obesity surgery typically experience changes in the levels of gut hormones, primarily glucagon-like peptide-1 (GLP-1) and peptide YY (PYY) (Chandarana, Gelegen et al. 2013, Guida, Stephen et al. 2019, Dischinger, Hasinger et al. 2020, De Bandt, Rives-Lange et al. 2022). Current pharmacotherapies, including GLP-1 receptor agonists (GLP-1RAs), primarily target a single receptor and signaling pathway. These receptor agonists have been used successfully to treat type 2 diabetes mellitus (T2DM), which is a frequent co-morbidity of obesity. However, the use of these drugs has been associated with several notable side effects, including malaise, nausea, and emesis, as well as other gastrointestinal issues. These drugs also have similar shortcomings when used to treat obesity alone (Borner, Tinsley et al. 2022). In response to these concerns, several novel dual or triple agonists have been created based on the structures of gut hormone agonists that are complementary to GLP-1, including glucagon, glucose-dependent insulinotropic polypeptide (GIP), and peptide YY_3-36_ (PYY_3-36_) (Talsania, Anini et al. 2005, Chepurny, Bonaccorso et al. 2018, Frias, Nauck et al. 2018, Kjaergaard, Salinas et al. 2019, Milliken, Elfers et al. 2021, Ostergaard, Paulsson et al. 2021, Battelino, Bergenstal et al. 2022, Boland, Laker et al. 2022, Heise, Mari et al. 2022, Jastreboff, Aronne et al. 2022, Metzner, Herzog et al. 2022, Zhao, Yan et al. 2022).

PYY_1–36_ is a gut hormone that binds to the Y1-R in pancreatic islets and central nervous system (CNS) nuclei that control appetite regulation in the brain including the brainstem area postrema (AP) and nucleus tractus solitarius (NTS), where it has an anorectic effect (Walther, Morl et al. 2011). The results of recent work reveal that GLP-1R is expressed in neuropeptide Y (NPY)-positive neurons in the AP and that GLP-1 can directly or indirectly inhibit neuronal signaling in the anorexigenic NPY system via agonism of GLP-1R (Ruska, Szilvasy-Szabo et al. 2022). Likewise, results from a considerable body of research revealed that PYY_1-36_ agonism of Y1-R plays a key role in promoting β-cell survival; this pathway has been recognized as critical to the reversal of diabetes and the recovery of impaired islet function after bariatric treatment (Guida, Stephen et al. 2017, Guida and Ramracheya 2020, Lafferty, Flatt et al. 2021). Acute administration of a Y1-R agonist may also reduce the rate of insulin secretion by pancreatic β-cells (Guida, Stephen et al. 2017, Guida, Stephen et al. 2019). Recent results demonstrating agonism at the arcuate nucleus (ARC) revealed that signaling via Y1-Rs protected female mice from obesity (Paterlini, Panelli et al. 2021, Oberto, Bertocchi et al. 2022). Other studies performed in mice demonstrate that pancreatic Y1-R activation facilitates the trans-differentiation of α-cells into β-cells, thereby improving insulin sensitivity (Lafferty, Flatt et al. 2021). The authors of this study noted that the pancreatic islets clearly benefited from Y1-R activation regardless of the diabetes status of the host (Lafferty, Flatt et al. 2021). Tanday *et al*., (2022) have since described a role for Y1-R agonism in inducing periods of β-cell rest that, when combined with GLP-1R agonism, proved beneficial in obesity-driven models of diabetes.

PYY_3–36_, a truncated peptide agonist derived from dipeptidyl peptidase IV (DPPIV)-mediated proteolysis of full-length PYY_1–36_, is of particular interest as it binds preferentially to the anorectic neuropeptide Y2-R (Talsania, Anini et al. 2005, Kjaergaard, Salinas et al. 2019, Merkel, Moreno et al. 2021, Ostergaard, Paulsson et al. 2021, Metzner, Herzog et al. 2022). PYY_3-36_ crosses the blood-brain-barrier (Nonaka, Shioda et al. 2003) and inhibits food intake via its interactions with Y2-R in brain areas that regulate energy homeostasis, including the ARC of the hypothalamus and the AP and NTS of the hindbrain (Shaw, Gackenheimer et al. 2003, Fetissov, Byrne et al. 2004, Neary, Small et al. 2005, Blevins, Chelikani et al. 2008). While circulating levels of PYY_3-36_ are frequently reduced in individuals with obesity (Batterham, Cohen et al. 2003, Roth, Enriori et al. 2005, Batterham, Heffron et al. 2006, Rahardjo, Huang et al. 2007, Roth, Bongiovanni et al. 2010), these levels typically return to those detected in average-weight individuals following a reduction in body weight and/or gastric bypass surgery (Roth, Enriori et al. 2005, Batterham, Heffron et al. 2006) (Roth, Bongiovanni et al. 2010).

Peripheral administration of PYY_3-36_ reduces caloric intake, increases postprandial insulin levels, and enhances insulin sensitivity, thermogenesis, lipolysis, and fat oxidation in both lean and obese humans as well as in nonhuman primates (Koegler, Enriori et al. 2005, Moran, Smedh et al. 2005) (Vrang, Madsen et al. 2006, Sloth, Davidsen et al. 2007, Abdel-Hamid, Abdalla et al. 2019). Administration of PYY_3-36_ also results in improved glucose control, improved lipid metabolism, and diminished insulin resistance in rodent models (Vrang, Madsen et al. 2006, van den Hoek, Heijboer et al. 2007, Chandarana, Gelegen et al. 2013). However, several significant limitations have hampered the development of PYY_3-36_ as an anti-obesity drug, including its short half-life (∼12 min) (Addison, Minnion et al. 2011) and its inability to sustain weight reduction beyond a 1–2-week period (Reidelberger, Haver et al. 2011). The inability to sustain weight loss may be due to PYY_3-36_-induced Y2-R down-regulation and tolerance (i.e., tachyphylaxis) and/or activation of compensatory mechanisms in response to reduced food intake.

Recent work suggested that the reduction in both food intake and body weight observed in response to combined treatment with GLP-1RAs and PYY_3-36_ results in synergistically-enhanced activation of discrete hypothalamic and brainstem circuits that regulate appetite, including the arcuate nucleus (ARC), the paraventricular nuclei of the hypothalamus (PVNs), the central nucleus of the amygdala (CeA), and the hindbrain (AP/NTS) (Dischinger, Corteville et al. 2019, Kjaergaard, Salinas et al. 2019, Lafferty, Flatt et al. 2021, Metzner, Herzog et al. 2022). Combination therapy has also resulted in improved glucoregulation via an increase in insulin sensitivity, an observation that has been linked to the recovery of pancreatic β-cell function (Lafferty, Flatt et al. 2021).

Based on previous research and promising clinical data (Guida, Stephen et al. 2017, Guida and Ramracheya 2020), we have designed a single chimeric peptide intended to target GLP-1R, Y1-R, and Y2-R-mediated pathways simultaneously. While other multi-agonists that target a variety of receptors have been explored, this specific approach has not been utilized in any previous studies. This approach offers a unique combination of potent weight-loss, glucoregulation with b-cell mass protection/proliferation, devoid of nausea/malaise.

To this end, we recently published the first description of GEP44, a chimeric peptide monomer that includes partial amino acid sequences of the GLP-1RA, Exendin-4 (Ex-4), and PYY. Results from our previous study revealed that GEP44 binds to GLP-1R, Y1-R, and Y2-R (Milliken, Elfers et al. 2021). Administration of GEP44 resulted in potent anorectic effects in two-to-five-day treatment studies in both lean and diet-induced obese (DIO) rats. Notably, administration of GEP44 resulted in an 80% reduction in food intake and a more profound loss of body weight than what could be achieved in response to treatment with several of the US Food and Drug Administration (FDA)-approved GLP-1RAs, including Ex-4 and liraglutide (LIRA). Furthermore, administration of GEP44 resulted in no visceral malaise, as determined in experiments with both rats and shrews (Milliken, Elfers et al. 2021).

In this manuscript, we develop the biochemical and preclinical aspects of GEP44. We found that GEP44 exhibits biased agonism at the Y2-R, as documented by its inability to induce Y2-R-mediated internalization. As a further development of the biased agonist approach, we introduce the GEP44 analog, GEP12. GEP12 was designed to incorporate findings published by Jones, Buenaventura et al. (2018), who reported that conversion of the N-terminal histidine (His) of Ex-4 to phenylalanine (Phe) resulted in reduced GLP-1R-mediated internalization and prolonged signal response. Interestingly, while administration of GEP12 did not result in a reduction in GLP-1R internalization, when evaluated in the presence of a Y1-R antagonist, it elicited increased insulin secretion in rat islets compared to results obtained with either GEP44 or Ex-4. These findings are consistent with the reduced rates of insulin secretion observed in islets in response to agonism at Y1-R (Yang, Ann-Onda et al. 2022).

Our findings highlight the potential of simultaneously targeting GLP-1R, Y1-R, and Y2-R and provide evidence suggesting a significant role for Y1-R agonism in the control of food intake and glucoregulation. This approach offers the possibility of achieving the long-term benefits of obesity surgery with a non-invasive approach. This will help millions of patients who are currently suffering from obesity and its co-morbidities, especially those for whom surgery is not an option.

## RESULTS

### GEP44 binds to and activates GLP-1R, Y1-R, and Y2-R via biased agonism

The addition of GEP44 to H188-GLP-1R transduced HEK293 cells resulted in elevated levels of cAMP with an EC_50_ value of 417 pM (**Fig. 1A**). The overall magnitude of response to GEP44 (and GEP12, *vide infra*) was nearly equivalent to the response to Ex-4 (**Fig. 1A**), but with differences noted in EC_50_ values for GEP44 (492.6 pM), GEP12 (17.3 nM) and Ex-4 (28.7 pM) at the GLP-1R (**Fig. 1A**). GEP44 agonism at Y1-R and Y2-R was validated in assays that monitored their ability to counteract adenosine-stimulated cAMP production in HEK293 C24 cells that were transiently transfected with Y1-R or Y2-R (Milliken, Doyle et al. 2020, Milliken, Elfers et al. 2021). As we reported previously (Milliken, Elfers et al. 2021), pre-treatment of these cells with GEP44 for 20 min resulted in a concentration-dependent inhibitory effect (**Fig.1B** and **1C**) with IC_50_ values of 34 nM and 27 nM for Y2-R and Y1-R, respectively. As shown in **Fig. 1D–F**, half-maximal GEP44 binding at GLP-1R was observed at 113 nM (*versus* 5.85 nM for Ex-4), 65.8 nM at Y2-R (*versus* 1.51 nM for PYY_3-36_), and 86.6 nM at Y1-R (*versus* 7.9 nM for PYY_1-36_).

**Figure 1.**
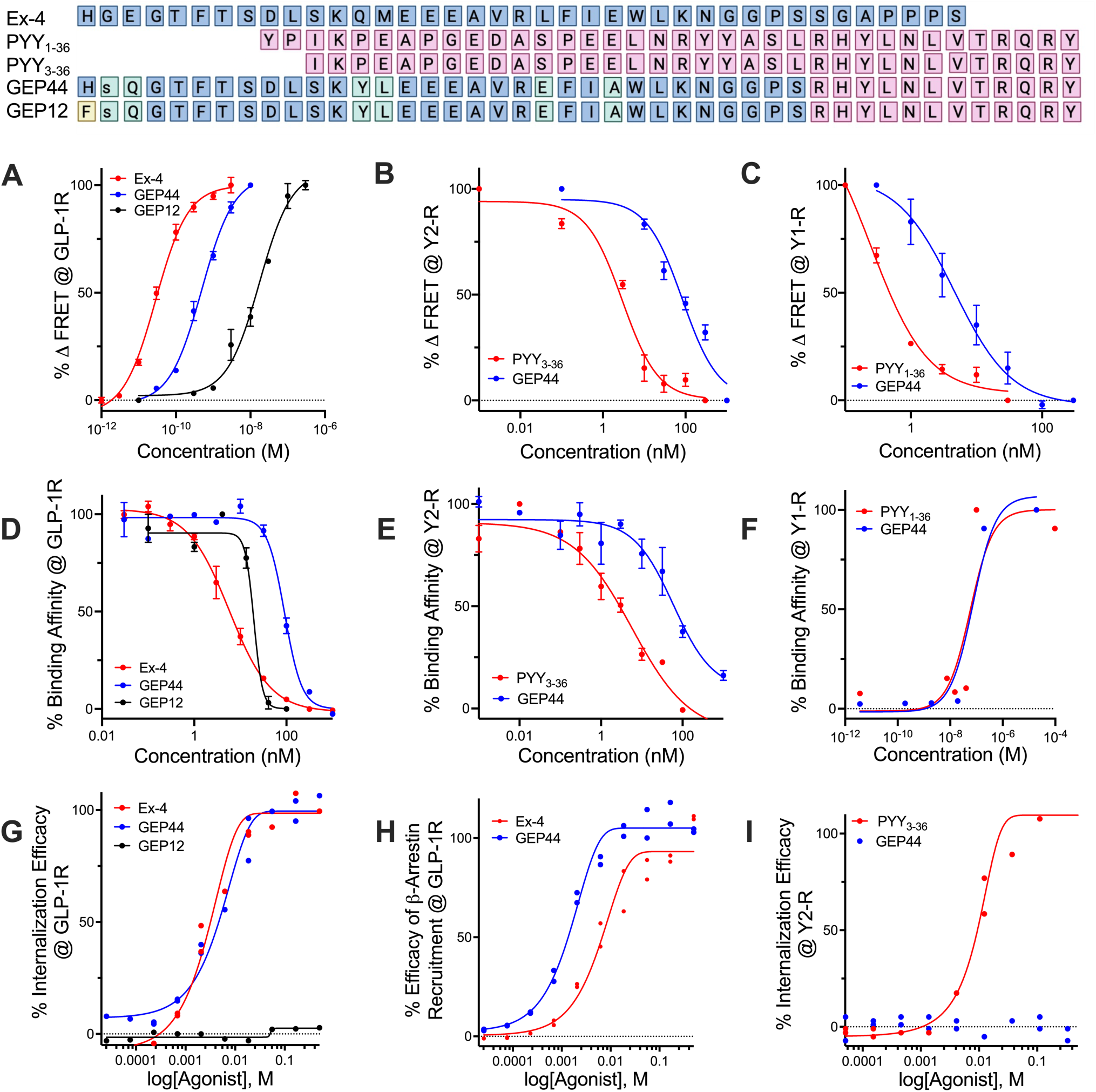
Design and *in vitro* evaluation of chimeric peptides GEP44 and GEP12. Shown are the amino acid sequences of Ex-4, PYY_1-36_ and PYY_3-36_ overlaid with those of GEP44 and GEP12 with lowercase single-letter amino acid code denoting a D-isomer. (**A**) Dose-dependent agonism (% change in FRET ratio tracking levels of cAMP) of Ex-4, GEP44, and GEP12 at the GLP-1R. (**B**) Dose-dependent agonism (% change in FRET ratio tracking levels of cAMP) of PYY_3-36_, GEP44, and GEP12 at the Y2-R. (**C**) Dose-dependent agonism of PYY_1-36_, GEP44, and GEP12 at the Y1-R. (**D**) Percent binding of Ex-4, GEP44, and GEP12 at the GLP-1R. (**E**) Percent binding of PYY_3-36_, GEP44, and GEP12 at the Y2-R. (**F)** Percent binding of PYY_1-36_, GEP44, and GEP12 at the Y1-R. (**G**) % internalization of GEP44 and GEP12 at the GLP-1R. (**H**) % recruitment of β-arrestin-2 by Ex-4 and GEP44 at the GLP-1R. (**I**) % internalization of Ex-4 and GEP44 at the Y2-R.

Biased agonism is a term used to describe ligand-mediated activation of a specific subset of the intracellular signaling pathways linked to a single receptor (Andresen 2011). With this in mind, we explored the GEP44-mediated internalization of Y2-R and internalization and recruitment of β-arrestin-2 at GLP-1R (**Fig. 1G–I**). Our results revealed that GEP44 did not promote internalization of Y2-R (**Fig. 1I**). This is notable given that Y2-R internalization is a critical step in receptor desensitization; this most likely results from strong allosteric effects associated with Y2-R/Gα interactions that place the cell in a refractory state and prevent further signaling (Ziffert, Kaiser et al. 2020).

By contrast, GEP44 elicited responses that were similar to those of the peptide agonist Ex-4 in assays designed to examine both internalization (**Fig. 1G**) and β-arrestin recruitment (**Fig. 1H**) at GLP-1R. Our findings revealed EC_50_ values of 3.97 nM and 2.43 nM for GLP-1R internalization mediated by GEP44 and Ex-4, respectively, and EC_50_s of 1.59 nM and 7.13 nM for GEP44 and Ex-4, respectively, for β-arrestin-2 recruitment.

A recent publication by Jones, Buenaventura et al. (Jones, Buenaventura et al. 2018), reported the impact of GLP-1R trafficking on insulin release and noted specifically that Ex-4 analogs with reduced capacity to elicit internalization and β-arrestin recruitment were more efficacious at inducing insulin release than the parent Ex-4 peptide. These results suggest that ligand-induced insulin secretion tracks inversely with β-arrestin recruitment. The authors reported that conversion of the N-terminal His to Phe resulted in an Ex-4 analog with improved efficacy at inducing insulin release while eliciting reduced levels of receptor internalization and β-arrestin-2 recruitment (Jones, Buenaventura et al. 2018). We used this information to create GEP12 (**Fig. 1**) which is a GEP44 analog with an analogous N-terminal His to Phe modification. We found that GEP12 functioned as a GLP-1R agonist with an EC_50_ of 17.3 nM (**Fig. 1A**) and bound to this receptor with an IC_50_ of 19.2 nM (**Fig. 1D**). As predicted, we observed no measurable GEP12-mediated internalization of GLP-1R in response to concentrations as high as 400 nM (**Fig. 1G**). Thus, we explored the responses to both GEP12 and GEP44 in *ex vivo* studies of insulin secretion in rat and human islets (**Fig. 2A** and **B**) as discussed in detail in the section to follow.

**Figure 2.**
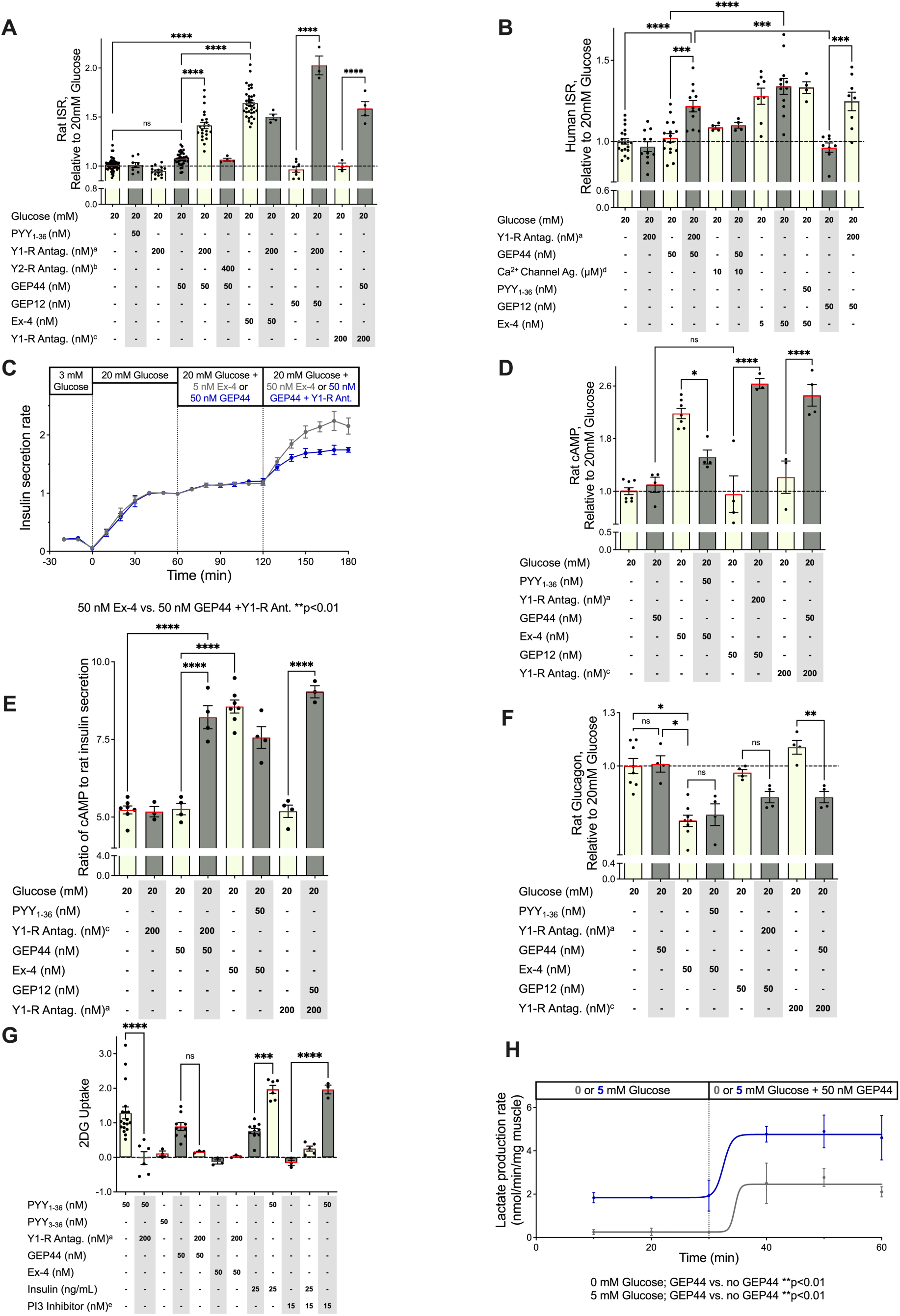
Action of GEP44 and GEP12 are mediated by GLP-1R and Y1-R in isolated pancreatic islets and muscle tissue. Rat (**A**) and human (**B**) islets were incubated for 60 min in 20 mM glucose and additional agents as indicated. Supernatants were then assayed for insulin, cAMP, and glucagon concentrations. (**C**) Insulin secretion rates (ISRs) were measured by perifusion over a one-hour incubation period in rat islets in 20 mM glucose with 5 or 50 nM peptides with or without Y1-R antagonist, as indicated (**C**). Impact of GEP44 on (**D**) cAMP, (**E**) the ISR to cAMP ratio, and (**F**) glucagon secretion, relative to glucose-mediated stimulation alone in the absence of test compounds. cAMP levels corresponded directly, and glucagon secretion corresponded inversely with the ISR. (**G**) Uptake of ^3^H-2-deoxyglucose (2-DG) and (H) lactate production (± 5 mM glucose) in response to GEP44 and other agents known to interact with GLP-1R, Y1-R, and Y2-R in the rat quadriceps muscle *ex vivo*. Horizontal dashed line in 2A, B, D, and F represents response to 20 mM glucose alone in assay as described. ^a^ = PD 160170; ^b^ = BIIE0246; ^c^ = BIBO; ^d^ = Bay K, ^e^ = Wortmannin.

### Effects of GEP44 and GEP12 on islets and muscle are mediated by both GLP-1R and Y1-R

Insulin secretion from rat and human islets was measured in the presence of 20 mM glucose and the presence or absence of various test compounds using both static and perifusion analyses. As shown in **Fig. 2A** and **B**, a static analysis revealed that both Ex-4 and GEP44 potentiated the insulin secretion rate (ISR) by 62% and 37%, respectively, over 20 mM glucose alone. However, a stimulatory response of GEP44 was observed only in the presence of Y1-R-specific antagonists (PD160170 or BIBO3304 - **Fig. 2A** and **B**), as expected if Y1-R agonism by GEP44 serves to counteract GLP-1R agonism by the same peptide. Our findings from perifusion experiments were consistent with those obtained from the static analysis. The results shown in **Fig. 2C** confirmed that GEP44 had a stimulatory effect on ISR in both rat and human islets and that this response occurred only in the presence of a Y1-R antagonist. The perifusion experiments also documented similar kinetic ISR responses to Ex-4 and GEP44 when islets were treated with a Y1-R antagonist (**Fig. 2C**). Nonetheless, the impact of GEP44 with a Y1-R antagonist in the doses used in this experiment is significant and greater than the responses observed to 5 nM Ex-4, which is the typical pharmacologic level of this agent (**Fig. 2B**). Interestingly, GEP12 elicited stronger increases in the ISR (**Fig. 2A** and **B**). Likewise, responses to both GEP44 and GEP12 were accompanied by significant increases in cAMP levels in both rat and human islets when measured the presence of glucose and Y1-R antagonists (**Fig. 2D** and **E).**

Taken together, the ISR and cAMP findings support the hypothesis that GLP-1R-mediated increases in cAMP potentiate insulin secretion, but that simultaneous occupation of Y1-R prevents binding to GLP-1R and/or the capacity to elicit relevant downstream responses. However, to explore the possibility that these peptides might modulate glucose homeostasis via their effects on circulating levels of glucagon, glucagon secretion from islets treated with GEP44 or GEP12 was measured, both in the presence and the absence of Y1-R antagonists (**Fig. 2F**). Our findings revealed that all agents and relevant combinations (i.e., GEP44 and GEP12 with Y1-R antagonists or Ex-4 alone) resulted in decreased glucagon release. Interestingly, this was the opposite of their individual impacts on the ISR (**Fig. 2F**). These data are consistent with one of two scenarios, either (1) direct peptide-stimulated inhibition of GLP-1R and Y1-R on α-cells, or (2) indirect peptide-mediated insulin release leading to inhibition of glucagon release, as observed in previous studies (Vergari, Knudsen et al. 2019); (Bansal and Wang 2008).

### GEP44 activates insulin-independent glucose uptake in muscle tissue via its interactions with Y1-R

Despite the positive ISR data and hypothesis that GEP12 would prove superior to GEP44, we noted in *in vivo* studies (*vide infra*) that it was considerably *inferior* to GEP44 in terms of food intake and body weight reduction. GEP12 is also poorly soluble and unstable (i.e., forms aggregates in solution). Based on these observations with GEP12, and on our unexpected findings that GEP44-mediated increases in ISR and decreases in glucagon release can be detected only in the presence of Y1-R antagonists, we proceeded to characterize the contributions of Y1-R to glucose uptake in muscle tissue. Uptake of the radiolabeled glucose analog, ^3^H-2-deoxyglucose (2-DG; **Fig. 2G**), was used to measure the impact of acute stimulation of glucose transport in freshly harvested rat quadriceps muscle. Our findings revealed that Y1-R signaling directly stimulated glucose uptake in muscle tissue via a pathway that is independent of insulin-stimulation. Addition of wortmannin, a biochemical inhibitor of AKT signaling that is central to insulin-mediated stimulation of glucose transport (Thiel, Guethlein et al. 2021) effectively blocked insulin-stimulated glucose uptake in muscle tissue. However, wortmannin had no impact on glucose uptake stimulated by the Y-1R agonist, PYY_1-36_ (**Fig. 2G**). Thus, based on the clear and direct role of Y1-R in promoting glucose uptake in muscle, the lack of any response to GLP-1R agonism, and the glucose-reducing effects of GEP44 observed in our previous studies (Milliken, Elfers et al. 2021), we predicted that GEP44 would also promote glucose uptake via interactions with the Y1-R. Accordingly, we performed parallel studies designed to measure the impact of PYY_1-36_, GEP44, and GEP44 together with the Y1-R-specific antagonist, PD160170, on glucose uptake into muscle (**Fig. 2G**). In contrast to the results of our previous experiments performed using pancreatic islets (i.e., those in which GEP44-mediated effects were more prominent in the presence of the Y1-R antagonist), we found that GEP44 directly increased glucose uptake in muscle tissue to a degree similar to that elicited by PYY_1-36_ and that GEP44-mediated glucose uptake was inhibited by Y1-R antagonism. To reinforce these findings, we also measured the release of lactate from muscle using a flow system (**Fig. 2H**). Based on our findings that revealed the ability of PYY_1-36_ to stimulate insulin-independent glucose transport, we further investigated the impact of this peptide and GEP44 on glycolysis. We reasoned that stimulation of glucose transport would have an impact on the rate of glycolysis as reflected by lactate release and if so, would provide a demonstration this novel effect of Y1R agonism on muscle based on the results of two different assay systems. Accordingly, we used the flow system that was featured previously in our experiments focused on islet analysis and observed a doubling of lactate production in response to GEP44 (**Fig. 2H**).

### Detection of GEP44 in Y1-R, Y2-R, and GLP-1R-expressing cells in caudal brainstem nuclei associated with appetite control (AP/NTS)

To visualize GEP44 localization in the brain, we injected rats with fluorescently labeled (f)-Cy5-GEP44 either intraperitoneally (IP; 15.5 μg/kg) or directly into the fourth ventricle (intracerebroventricular injection [ICVI]; 1 μg/μl) to target caudal brainstem regions (e.g., the AP and the NTS) involved in food intake and control of nausea/malaise. To evaluate f-Cy5-GEP44 localization in Y1-R, Y2-R, and/or GLP-1R-expressing cells in the rat hindbrain, we performed RNAscope fluorescent *in situ* hybridization (FISH) as previously described (Fortin, Lipsky et al. 2020) combined with immunohistochemistry (IHC) of coronal sections prepared from the rat brainstem. f-Cy5-GEP44 administered IP (**Fig. 3A**) or by ICVI (**Fig. 3B**) was detected in the brainstem at 60 min post-injection, specifically in Y1-R and/or GLP-1R-expressing cells within the AP and the NTS. We also detected f-Cy5-GEP44 in Y1-R, Y2-R, and GLP-1R-expressing cells in these regions (**Fig. 3C**). Localization of f-Cy5-GEP44 in cells expressing Y1-R or Y2-R was observed primarily within the medial NTS, while GEP44 localization in cells expressing GLP-1R was detected primarily within the AP.

**Figure 3.**
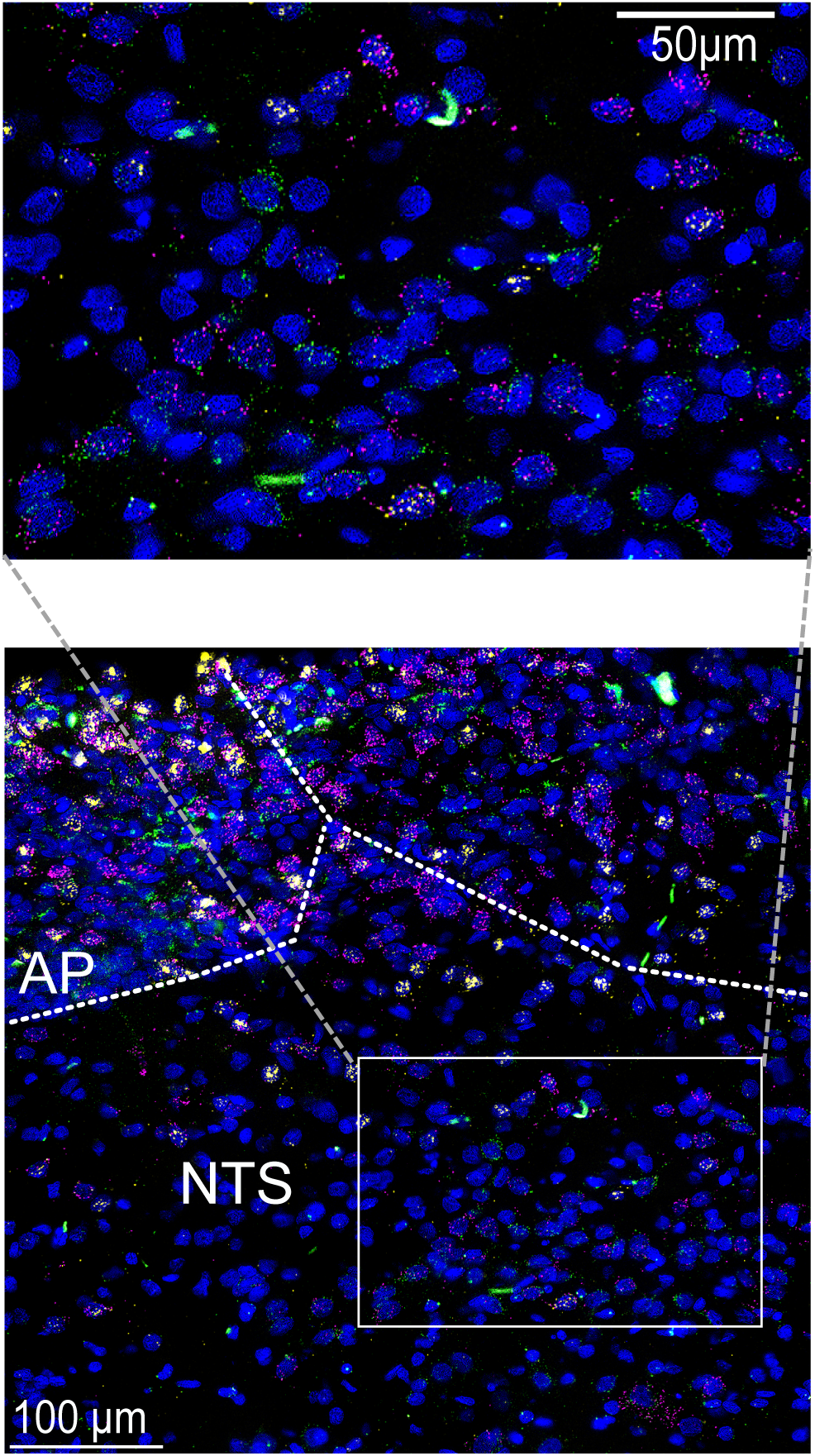
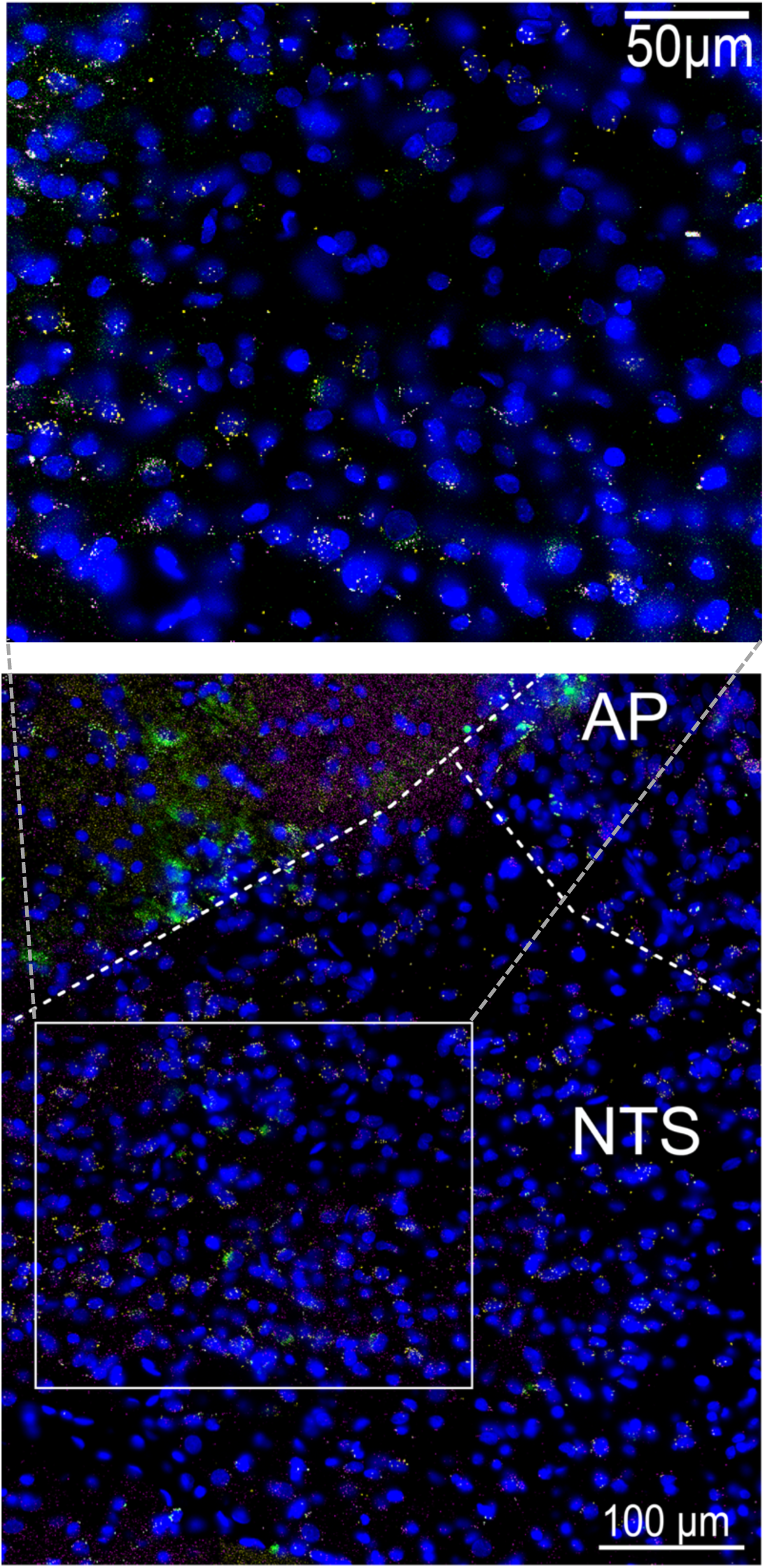
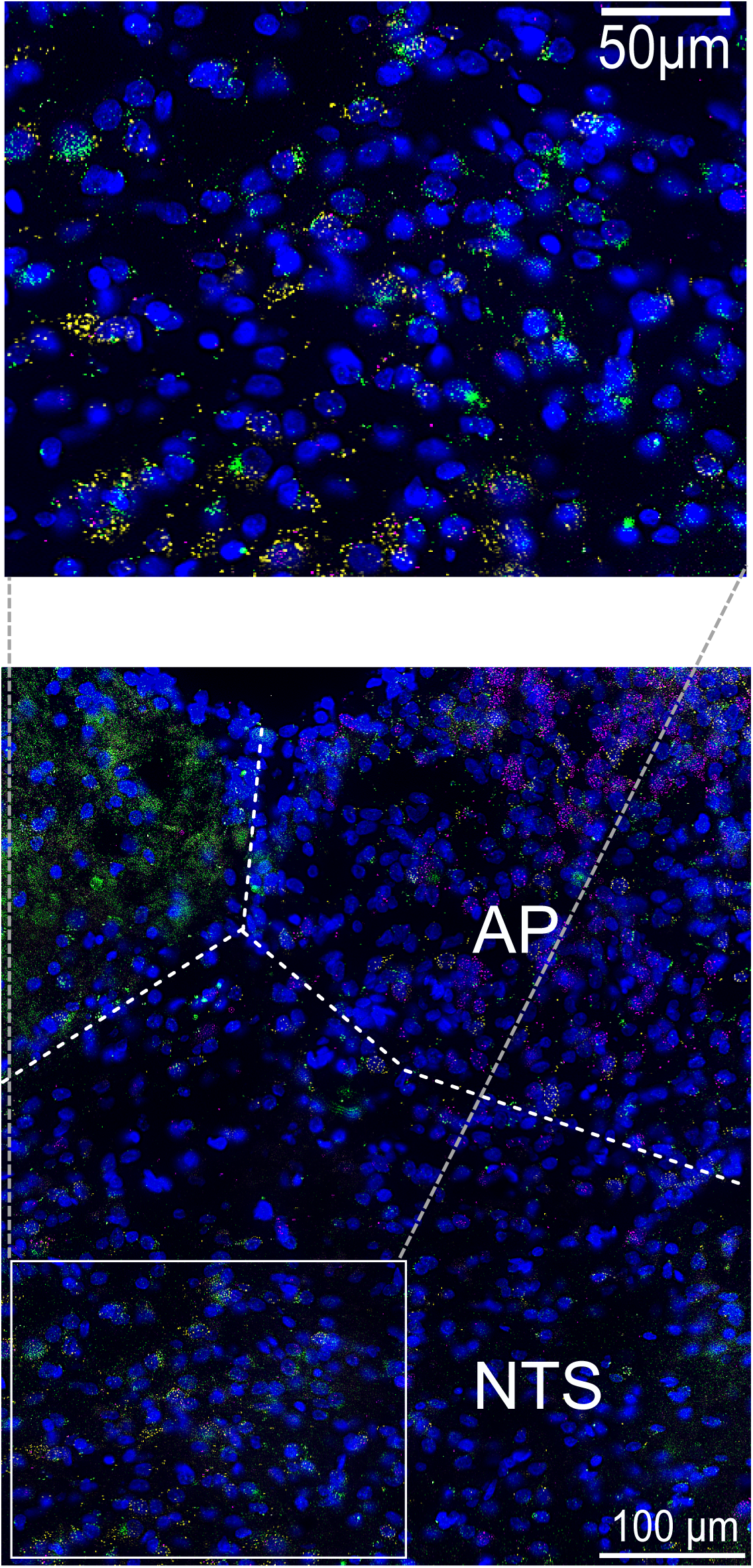
FISH and IHC visualization of f-Cy5-GEP44 and its colocalization with Y1-R, Y2-R, and GLP-1R in cells in the NTS/AP regions of the rat brain. (A) f-Cy5-GEP44 (green) administered IP colocalized with Y1-R and GLP-1R (yellow) in the AP. (B) f-Cy5-GEP44 administered ICVI colocalized with Y1-R (yellow) and GLP-1R (magenta) in the AP. See Supplementary Video 1 for a three-dimensional (3D) rotational image of the area within the inset. (C) f-Cy5-GEP44 administered ICVI colocalized with Y2-R (yellow) and GLP-1R (magenta) in cells of the AP. See Supplementary Video 2 for a 3D rotational image of the area within the inset. Images are shown at 40x magnification.

### *In vivo* dose-response on food intake/body weight in diet-induced obese (DIO) rats

In our first preclinical experiment, we performed a dose escalation study in DIO rats to compare responses to treatment with GEP44 and Ex-4. Our results revealed similar reductions in food intake in response to daily injections of GEP44 or Ex-4 at concentrations ranging from 0.5–10 nmol/kg. However, rats treated with GEP44 at 20 nmol/kg/day exhibited greater reductions in food intake compared to those treated with the same concentration of Ex-4 (predicted mean difference, -18.5%; 95% confidence interval [CI], -36.3% to -0.7%, *p*=0.0382). Notably, rats receiving Ex-4 at doses of 5 nmol/kg and higher exhibited a lack of activity during the first 30 min of the dark cycle relative to rats injected with vehicle alone. With doses of 10 nmol/kg and higher, rats exhibited not only a lack of activity, but also notable changes in facial expressions associated with pain and nausea including orbital tightening and nose/cheek flattening compared to rats treated with vehicle alone. Thus, we limited the doses of Ex-4 used in these experiments to ≤20 nmol/kg/day. By contrast, we were able to continue the dose escalation in rats treated with GEP44. Our findings revealed that rats receiving 50 or 100 nmol/kg/day responded with ∼77% and ∼90% reductions in food intake relative to baseline, respectively (**Fig. 4A**). The minimal effective dose (MED) determined for both drugs was 0.5 nmol/kg/day. Our results indicated a ‘no observed adverse effect level’ (NOAEL) of 2 nmol/kg/day for Ex-4 and a maximal tolerated dose (MTD) of 20 nmol/kg/day. By contrast, the DIO rats tolerated much higher doses (up to 100 nmol/kg/day) of GEP44 with no indication of malaise. We did detect mild staining around the mouth and nose of one rat treated with 50 nmol/kg/day GEP44 and one additional rat when treated with a single 100 nmol/kg dose of GEP44. Staining did not appear to be the result of inattention to grooming and was not consistent with porphyrin secretion. All animals remained alert and responsive with no other notable changes in appearance, activity, or behavior compared to rats treated with vehicle alone. Thus, we established a NOAEL for GEP44 at 20 nmol/kg/day. No MTD was determined.

**Figure 4.**
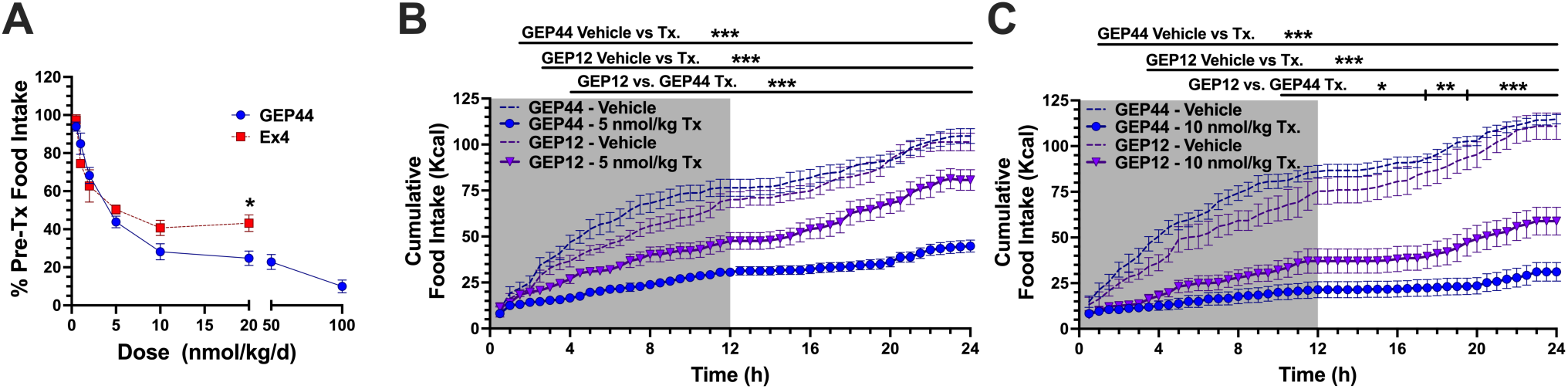
Dose-escalation study reveals a robust reduction of food intake in response to GEP44. (**A**) The dose-escalation study shows a robust reduction of food intake in response to GEP44 (●, n=8 DIO rats) *vs.* Ex-4 (▪, n=4 DIO rats). Food intake was averaged over three days of treatment at each drug dose and was normalized to the earliest three days during which all animals received injections with the vehicle control. Escalation of the Ex-4 dose was stopped at 20 nmol/kg due to multiple indicators of malaise. (**B, C**) Shown is the average 24-hour cumulative food intake for the three-day vehicle-treated baseline and three-day GEP12 treatment phases (▼, n=8 DIO rats) at (**B**) 5 nmol/kg/day and (**C**) 10 nmol/kg/day doses. Data from equivalent dose-testing performed as part of the GEP44 (●) dose-escalation study were included in these figures to facilitate a qualitative comparison. Data shown are means ± standard error of the mean (SEM); \**p*<0.05, ** *p* <0.01, *** *p* <0.001.

In additional experiments, treatment of DIO rats with GEP12 at 10 nmol/kg/day resulted in significant reductions in food intake and body weight (food intake: mean difference -41.3 kcal/day, 95% CI -62.8 to -19.9 kcal/day, *p*=0.0019; body weight: mean difference -32.6 g, 95% CI -53.5 to -11.7 g, *p*=0.0062). By contrast, no significant differences were observed in response to GEP12 at 5 nmol/kg/day (food intake: mean difference -20.0 kcal/day, 95% CI -41.4 to 1.5 kcal/day, *p*=0.067; body weight: mean difference -8.7 g, 95% CI -29.6 to 12.24 g, *p*=0.479). The impact of GEP12 on food intake relative to baseline was notably less than that observed in response to GEP44 (at 5 nmol/kg/day: mean difference 36.5%, 95% CI 14.9% to 58.0%, *p*=0.0009; at 10 nmol/kg/d: mean difference 34.6%, 95% CI -13.0% to 56.1%, *p*=0.0015). An analysis of 24-h food intake patterns revealed the GEP12-mediated suppression of food intake was of substantially shorter duration than that observed in response to GEP44 (**Fig. 4B** and **C**). No appreciable kaolin consumption was recorded.

### Long-term studies of GEP44 to treat diet-induced obesity *in vivo*

Our first longer-term experiment was a vehicle-controlled study designed to examine the responses of DIO rats to equimolar doses of GEP44 and liraglutide (LIRA) beginning with 10 nmol/kg/day and increasing to 25 nmol/kg/day after day 10. The rats treated with GEP44 exhibited notably larger reductions in both body weight (day 10: predicted mean difference -4.9%, 95% CI -6.6 to -3.3%, *p*<0.0001; day 16: predicted mean difference -7.9%, 95% CI -9.7 to -6.1%, *p* <0.0001) and food intake (10 nmol/kg/day: predicted mean difference -29.8 kcal/day, 95% CI -45.4 to -14.3 kcal/day, *p*=0.0001; 25 nmol/kg/day, predicted mean difference -34.1 kcal/day, 95% CI -49.7 to -18.6 kcal/day, *p* <0.0001) compared to rats treated with LIRA. Treatment with LIRA had no impact on body weight or food intake when administered at 10 nmol/kg/day and induced only mild transient changes at 25 nmol/kg/day (**Fig. 5A** and **B**). At the 25 nmol/kg/day dose, food intake was reduced by 62% in rats treated with GEP44 compared to 25% in rats treated with LIRA. At the end of the 16-day treatment protocol, we found that rats that received 25 nmol/kg/day exhibited an average body weight reduction of -12.1% compared to -3.2% in rats treated with LIRA. Blood glucose levels (**Table 1**) were lower on average in DIO rats treated with GEP44 compared to those treated with vehicle alone (estimated treatment effect, -8.1 mg/dL, 95% CI -15.3 to -1.0 mg/dL, *p*=0.026), while no difference was noted in LIRA vs. vehicle treated animals (estimated treatment effect, -3.4 mg/dL, 95% CI -10.2 to 3.3 mg/dL, p=0.309).

**Figure 5.**
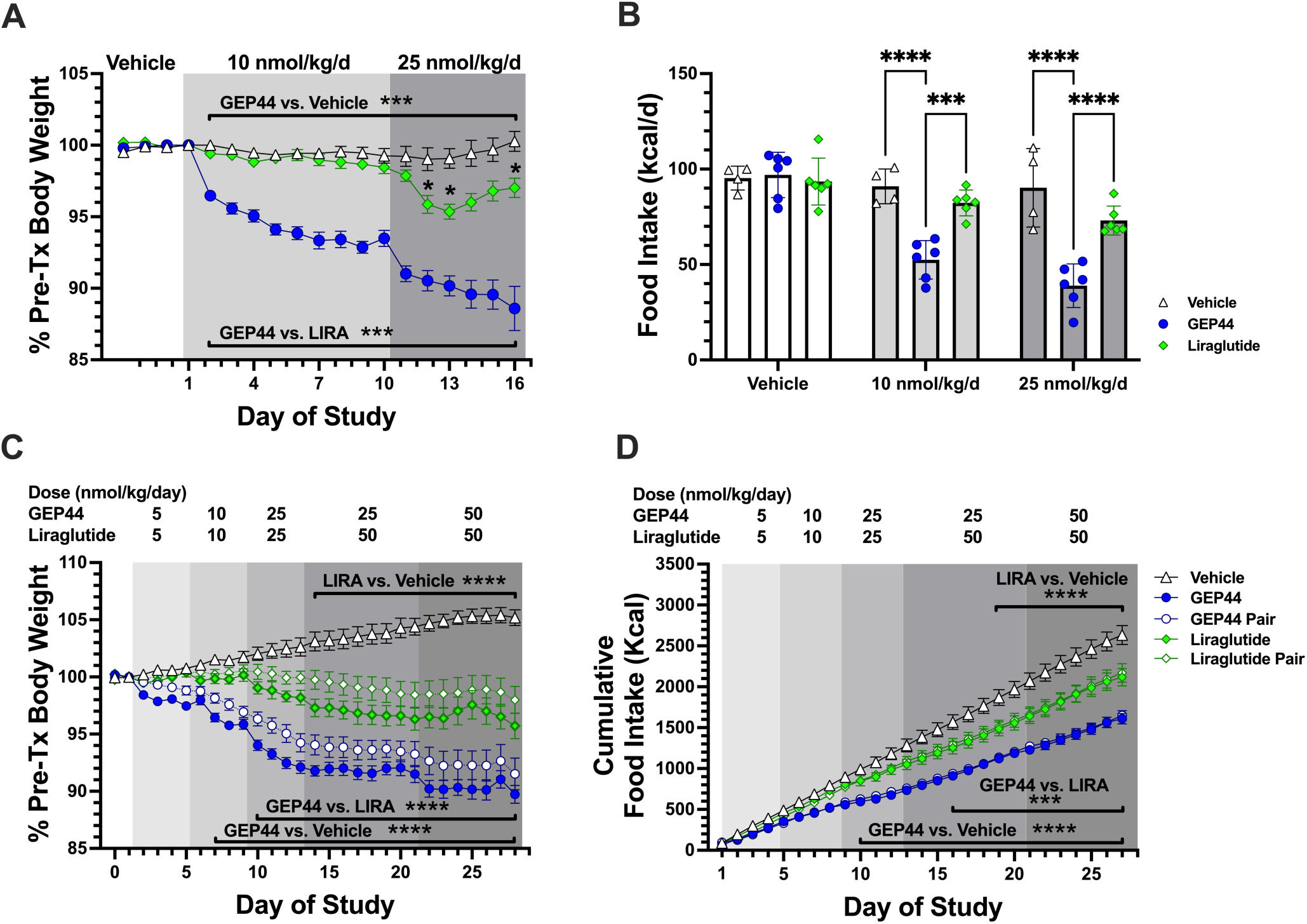
GEP44-mediated reductions in body weight and food intake were stronger than those elicited by LIRA during a 16-day and 27-day dose escalation protocol. (A, B) DIO Wistar rats were treated with vehicle (L) or with GEP44 (●) or LIRA (♦) at 10 nmol/kg/day for 9 days followed by 25 nmol/kg/day for 7 days (n=4–6) rats/group. In a second experiment, (C) changes in body weight and (D) food intake was evaluated during 27 days of treatment with vehicle, GEP44, vehicle-treated rats that were pair-fed to those receiving GEP44, LIRA, and vehicle-treated rats that were pair-fed to those receiving LIRA; n=8 per group. DIO male Wistar rats were matched based on baseline food intake and initial body weight gain trajectory. Changes in body weight were evaluated in response to GEP44 (●) at doses escalating from 5 to 50 nmol/kg/day. Rats underwent pair-feeding to match the amount of food consumed by their GEP44-treated counterparts (o). Other groups included rats treated with saline-vehicle control (△), LIRA (♦), and rats that were pair-fed to their LIRA-treated counterparts (◊). Symbols representing the results from pair-fed animals are overlayed by those from the GEP44 and LIRA treatment groups. Data shown are means ± SEM; \**p* <0.05, \*\*\**p* <0.001, \*\*\*\**p* <0.0001.

**Table 1.** Characteristics of treatment groups at baseline and after 27 days of treatment with GEP44 or LIRA. The data shown are means ± standard deviation (SD). Data were compared by ANCOVA followed by pairwise comparisons of marginal linear predictions using a Bonferroni correction; n=8 rats per group. Abbreviations: Tx, treatment; \**p* <0.05, \*\**p* <0.01, \*\*\**p* <0.001 *vs.* vehicle control; #*p* <0.05, ^##^*p* <0.01, ^###^*p* <0.001 *vs.* LIRA. Pair-fed comparisons were made with no difference detected.

|  | Vehicle | GEP44 |  | GEP44<br>pair-fed |  | Liraglutide |  | Liraglutide<br>pair-fed |  |
| --- | --- | --- | --- | --- | --- | --- | --- | --- | --- |
| **Body Weight (g)** |  |  |  |  |  |  |  |  |  |
| Baseline | 660 ± 94 | 660 ± 55 |  | 667 ± 63 |  | 660 ± 65 |  | 658 ± 75 |  |
| Post-Tx | 695 ± 104 | 592 ± 55 | *** ## | 611 ± 70 | *** | 631 ± 79 | *** | 647 ± 86 | *** |
| **Food Intake** |  |  |  |  |  |  |  |  |  |
| 5d Baseline (kcal/d) | 88.6 ± 12.6 | 95.6 ± 8.9 |  | 94.2 ± 14.4 |  | 93.1 ± 8.7 |  | 91.3 ± 4.3 |  |
| Last 5d of Tx (kcal/d) | 91.8 ± 10.4 | 65.5 ± 10.0 | *** ## | 65.6 ± 10.2 | *** | 78.3 ± 11.1 | *** | 78.5 ± 10.6 | * |
| Cumulative (kcal) | 2633 ± 329 | 1610 ± 184 | *** ## | 1637 ± 186 | *** | 2116 ± 300 | ** | 2166 ± 327 | * |
| **Glucose (mg/dL)** |  |  |  |  |  |  |  |  |  |
| Baseline | 120 ± 10 | 110 ± 8 |  | 119 ± 13 |  | 119 ± 10 |  | 119 ± 5 |  |
| Post-Tx | 93 ± 8 | 83 ± 6 | * | 90 ± 6 |  | 89 ± 3 |  | 93 ± 9 |  |
| **Insulin (ng/mL)** |  |  |  |  |  |  |  |  |  |
| Baseline | 14 ± 10 | 13 ± 5 |  | 16 ± 6 |  | 14 ± 5 |  | 14 ± 4 |  |
| Post-Tx | 14 ± 8 | 7 ± 2 | ** | 6 ± 3 |  | 8 ± 3 | * | 8 ± 3 |  |

The second long-term study focused on efforts to sustain these effects via a gradual dose escalation from 5 to 50 nmol/kg/day. The results of this study recapitulated the larger reductions in body weight (mean difference -5.8%, 95% CI -7.9 to -3.7%, *p*<0.0001) and food intake (mean difference -309 kcal, 95% CI -549 to -70 kcal, *p*=0.004) observed in response to GEP44 compared to equivalent doses of LIRA during the first 12 days of treatment (**Fig. 5C** and **D**). During the remaining 15 days, GEP44 treatment continued to yield greater reductions in food intake relative to baseline (mean difference -14.3%, 95% CI -25.4 to -3.3%, *p*=0.01), even though the rats were treated with stronger doses of LIRA on days 13 through 20. At the end of the 27-day treatment protocol, the observed reductions in body weight compared to the vehicle were -15% and -9% for rats treated with GEP44 and LIRA, respectively. Similarly, the cumulative reduction in food intake reduction was -39% for rats treated with GEP44 *versus* -20% for rats treated with LIRA. At the end of the 27-d treatment, body weight reduction compared to vehicle was -15% (GEP44) vs. -9% (LIRA), and cumulative food intake reduction was -39% (GEP44) vs. -20% (LIRA) (**Table 1**).

Finally, while reductions of mean body weight were consistently less for vehicle-treated rats that were pair-fed with those treated GEP44 or LIRA compared to their peptide-treated counterparts, the differences did not achieve statistical significance. Rats receiving either GEP44 or LIRA exhibited lower post-treatment fasting insulin levels compared to those treated with vehicle alone. Interestingly, vehicle-treated rats that were pair-fed to those receiving GEP44 exhibited comparatively lower fasting serum cholesterol and HDL levels than did their peptide-treated counterparts. No differences in fasting blood glucose, triglycerides, or hepatic transaminase levels were observed (**Table 2**).

**Table 2.**
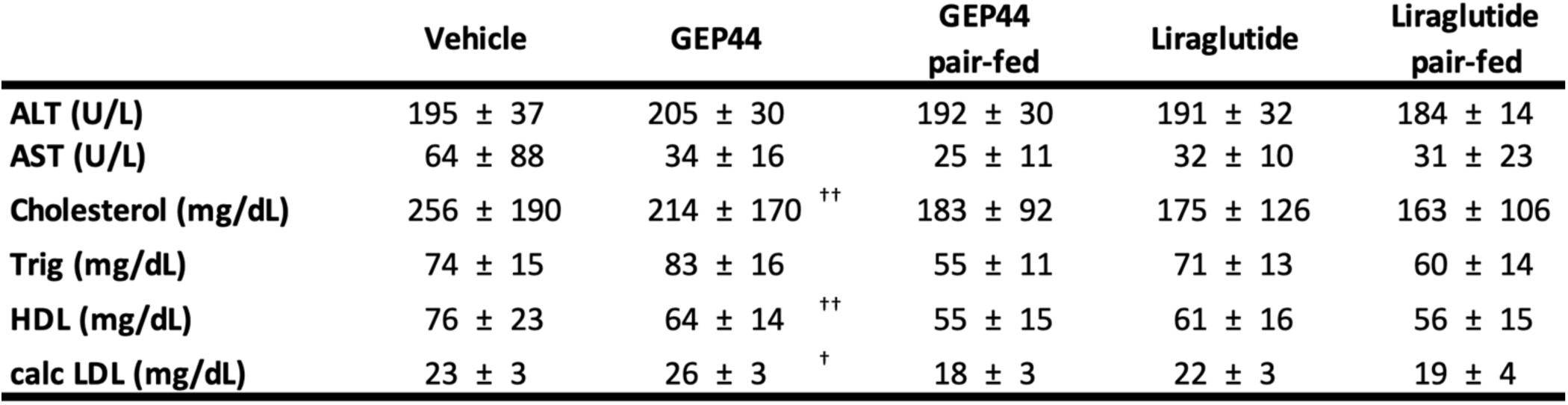
Outcomes after 27 days of treatment with GEP44 or LIRA. Data shown are means ± SD. Cross-sectional analyses were performed using an ANOVA followed by pairwise comparisons of means using a Bonferroni correction; n=8 rats per group. Abbreviations: ALT, alanine aminotransferase; AST, aspartate aminotransferase; Trig, triglycerides; HDL, high-density lipoprotein; calc, calculated; LDL, low-density lipoprotein; ^†^ *p* <0.05, ^††^ <0.01 *vs.* pair-fed counterparts.

## DISCUSSION

In this manuscript, we present the properties of the unimolecular triple agonist peptide, GEP44. GEP44 interacts with GLP-1, Y1, and Y2 receptors to regulate insulin secretion in both rat and human pancreatic islets, and it promotes insulin-independent Y1-R-mediated glucose uptake in rat muscle tissue *ex vivo.* Furthermore, the administration of GEP44 results in profound reductions in food intake and body weight in DIO rats. We modified GEP44 at its N-terminus to generate GEP12. As hypothesized, this modification resulted in reduced internalization at the GLP-1R. However, while there is a stronger stimulation of insulin secretion in response to GEP12 vs. GEP44, GEP12 it is less potent than GEP44 when administered *in vivo* to reduce food intake and body weight.

There are many seemingly conflicting reports on the effects of Y1-R agonism *versus* antagonism found in the recent literature. While receptor levels, animal diet, dosing, and the presence of a GLP-1RA have a profound impact on insulin secretion and associated endocrine outcomes, the effects of dual or triple agonism of Y1-R and Y2-R, in the presence of a GLP-1RA remain unclear. Several groups have designed monomeric dual and triple agonists based on the interactions of GLP-1 with glucagon (Day, Gelfanov et al. 2012, Sanchez-Garrido, Brandt et al. 2017, Ambery, Parker et al. 2018) and/or GIP (Finan, Yang et al. 2015, Nowak, Nowak et al. 2022, Tan, Akindehin et al. 2022). This is a novel and promising approach to the development of drugs for the treatment of human obesity. However, many of these drugs are poorly tolerated and are associated with significant adverse events. For example, once weekly dosing of tirzepatide (a dual GIPR and GLP-1R co-agonist) has superior efficacy for weight reduction compared to the GLP-1R agonist, semaglutide; at least 50% of patients receiving 10 or 15 mg of tirzepatide per week achieved a weight loss of 20% and more (Willard, Douros et al. 2020, Min and Bain 2021, Rosenstock, Wysham et al. 2021, Jastreboff, Aronne et al. 2022, Nowak et al. 2022).Tirzepatide was also superior to semaglutide at reducing levels of glycated hemoglobin (Frias, Davies et al. 2021). However, tirzepatide treatment resulted in mild to moderate gastrointestinal symptoms, including nausea (12–24%), diarrhea (15–17%) and vomiting (6–10%). These symptoms led to the discontinuation of treatment in 3-7% of those patients (Jastreboff, Aronne et al. 2022). Thus, our goal was to design a peptide agonist that overcomes these shortcomings and compare glucoregulation/appetite control driven by the incorporation of Y1-R/Y2-R agonism, a property not shared by tirzepatide incorporation. GEP44 was developed based on extensive structure-activity relationship studies and earlier preliminary *in vivo* studies that demonstrated its greater efficacy at reducing both food intake and body weight compared to the agonist peptide, Ex-4, without triggering gastrointestinal distress (as assessed by kaolin intake and behavioral scoring as a proxy for nausea in rats and direct evidence of emesis in shrews) (Milliken, Elfers et al. 2021).

Despite reports of positive outcomes from simultaneous agonism of both GLP-1R and Y2-R, this approach alone has not generated the same clinical benefits as obesity surgery (Ye, Hao et al. 2014, Boland, Mumphrey et al. 2019, Dischinger, Heckel et al. 2021). In one study, exogenous administration of combined GLP-1R/Y2-R agonists partially mimicked the positive effects of bariatric surgery but did not lead to the anticipated overall metabolic improvements (Metzner, Herzog et al. 2022) suggesting other pathways are also involved. Results of a study published by Dischinger, Hasinger et al. (2020) support this concept. Specifically, while obesity surgery and LIRA/PYY_3-36_ co-administration resulted in comparable changes in body weight in obese rats, only surgery resulted in profound changes to the hypothalamic transcriptome; these findings may explain in part the limited nature of the metabolic improvements observed in response to pharmacologic intervention (Dischinger, Hasinger et al. 2020, Metzner, Herzog et al. 2022). Others have demonstrated that the positive outcomes of obesity surgery (i.e., changes in body weight and improved glucose homeostasis) are sustained in mice in the absence of GLP-1R or Y2-R, or both GLP-1R and Y2-R (Ye, Hao et al. 2014, Boland, Mumphrey et al. 2019). These findings support the hypothesis that numerous intersecting and redundant pathways are involved in these physiologic responses. Bearing in mind the observed agonism and binding across GLP-1R, Y1-R and Y2-R (**Fig. 1**), we also determined that GEP44 delivered either peripherally or directly via ICVI localizes in hindbrain regions involved in the control of food intake. GEP44 was detected in AP/NTS cells that express GLP-1R, Y1-R, or Y2-R (**Fig. 3**). This result highlights a potential mechanism used by GEP44 to suppress food intake and promote reduction in body weight.

### Binding, agonism, internalization, and β-arrestin recruitment

Studies published by (Jones, Buenaventura et al. 2018) and (Ziffert, Kaiser et al. 2020) have documented the physiologic relevance of internalization and β-arrestin recruitment in response to specific interactions with both the GLP-1R and Y2-R. (Jones, Buenaventura et al. 2018) demonstrated that the extent of internalization and receptor trafficking mediated by Ex-4 (the scaffold upon which GEP44 is built) has a direct impact on the ISR and that efforts to retain GLP-1R on the cell surface result in enhanced and sustained insulin release. Similarly, (Ziffert, Kaiser et al. 2020) reported that Y2-R internalization results in receptor desensitization accompanied by a G_i_-refractory state. Thus, we evaluated GEP44-mediated GLP-1R and Y2-R internalization and compared the results to those obtained using Ex-4 or PYY_3-36_, respectively. Interestingly, GEP44 and Ex-4 were similarly effective at promoting GLP-1R internalization, with EC_50_ values of 3.97 nM and 2.43 nM, respectively. By contrast, while PYY_3-36_ exhibited the anticipated efficacy at Y2-R in this assay (IC_50_, 8.83 nM), GEP44 was not internalized. This unanticipated outcome may provide critical clues to the mechanism underlying GEP44-mediated weight loss observed in experiments performed *in vivo*. Thus, we modified GEP44 with the intent of *also* reducing its capacity to induce internalization at the GLP-1R. We hypothesized that this modified peptide would elicit improved ISRs in both rat and human islets. Based on information published (Jones, Buenaventura et al. 2018), we synthesized GEP12, a peptide with a single N-terminal amino acid change (His to Phe) from GEP44 (**Fig. 1**). GEP12 (IC_50_ = 19.2 nM) exhibited >4-fold greater binding affinity at human GLP-1R compared to GEP44 (IC_50_ = 90.4 nM) and elicited an increased ISR compared to either GEP44 (in the presence of a Y1-R antagonist) or Ex-4 alone. The N-terminal Phe residue in GEP12 likely contributes to its increased affinity for the extracellular binding domain of GLP-1R. As predicted, GEP12 elicits little to no GLP-1R internalization, a finding that is consistent with and confirms the observations of (Jones, Buenaventura et al. 2018), and is consistent with the concept of biased agonism. Biased agonism may be a critical factor underlying the observed effects of GEP44 and its GEP12 analog; similar results have been reported in studies that characterized the recent FDA-approved single molecule tirzepatide. These studies revealed that tirzepatide is an imbalanced and biased dual GIP-R/GLP-1R co-agonist, which stimulates GIP receptors in a manner analogous to its parent peptide but favors cAMP generation over β-arrestin recruitment at the GLP-1R. Thus, tirzepatide promotes enhanced insulin secretion when compared with responses to native GLP-1 (Willard, Douros et al. 2020).

### Y1-R-mediated effects on pancreatic islets

Our experiments were also designed to determine whether peptides that are efficacious at lower concentrations and/or those that promote modified internalization/β-arrestin recruitment might be capable of sustaining the beneficial effects beyond those currently observed while also reducing the frequency of adverse events. These improvements are likely to increase patient compliance and improve their overall quality of life. Our results suggest that the responses to GEP44 are most likely mediated via integrated responses from several tissues. We show in this manuscript that Y1-R signaling facilitates insulin-independent glucose uptake in muscle. This finding complements findings from an earlier study that revealed a role for Y1-R agonism in facilitating trans-differentiation of α-cells into β-cells (Lafferty, Flatt et al. 2021).

Initially, we anticipated that GEP44 would reduce serum glucose levels via binding to GLP-1R. Of note, we found that Ex-4 binding to GLP-1R leads to increased cAMP levels, an elevated ISR, and reduced glucagon secretion, all consistent with previous findings. However, our experimental studies with GEP44 revealed that interactions with Y1-R could mask the effects of GLP-1R agonism in isolated pancreatic islets. We found that GEP44 had little to no impact on glucose-stimulated insulin secretion in isolated islets in the absence of Y1-R antagonists. These findings may result from a direct interaction between Y1-R and GLP-1R and/or the impact of Y1-R signaling on GLP-1R-mediated induction of cAMP; the latter explanation is suggested by our results (see Fig. 2). The absence of GEP44-mediated changes to the ISR in the absence of a Y1-R antagonist suggests two possible mechanisms that would be consistent with the serum glucose-lowering effects observed in response to this chimeric peptide *in vivo*. First, we considered the possibility that responses mediated by GLP-1R might be modulated by the state of the Y1-R. In support of this hypothesis, we found that the Y1-R ligand, PYY_1-36_, inhibited Ex-4-mediated stimulatory responses. We also observed a direct correlation between cAMP levels and the ISR. Conversely, we also note the inverse relationship between glucagon release and the ISR. It is not yet clear whether this is the direct result of insulin-mediated inhibition of glucagon secretion or inhibition resulting from the activation of an α-cell receptor.

Our data suggest a role for Y1-R on pancreatic β-cell function, a role also suggested by other recent results (Yan, Zeng et al. 2021) documenting up-regulation of Y1-R mRNA in brown adipose tissue, inguinal white adipose tissue, and skeletal muscle of obese mice and humans. Other studies have documented up-regulation of the Y1-R agonist, PYY_1-36_, in pancreatic islets after obesity surgery; this ligand is then converted to the Y2-R agonist PYY_3-36_ via enzymatic cleavage by DPPIV (Chan, Mun et al. 2006); (Ballantyne 2006). Thus, DPPIV inhibitors may function by prolonging the half-lives of both GLP-1 and PYY_1-36_ *in vivo* (Aaboe, Knop et al. 2010). A 12-week study in which patients diagnosed with T2DM were treated with the DPPIV inhibitor, sitagliptin, revealed increased serum levels of PYY_1-36_ and improvement of glucose and non-glucose dependent insulin secretion (Aaboe, Knop et al. 2010). Therefore, DPPIV inhibitors might promote glucoregulation in part because of resulting elevated levels of PYY_1-36_, albeit with associated losses in food intake and body weight reduction.

### GEP44 mediates insulin-independent glucose uptake in muscle via interactions with Y1-R

GEP44 may also reduce serum glucose levels via direct interactions resulting in increased glucose uptake in muscle tissue. Given the known role of Y1-R in GLP-1R-mediated regulation of insulin secretion, we were surprised to find that Y1-R also stimulated glucose uptake in muscle via a mechanism that was fully independent of insulin stimulation. GEP44-mediated increases in glucose transport also correlated with its stimulation of lactate production and thus a downstream impact on muscle metabolism; however, Y1-R agonism may also stimulate a reaction step in the glycolytic pathway. Given that previous findings suggested a role for insulin-independent glucose uptake in pathways leading to glucose homeostasis (Wiernsperger 2005, Diener, Mowbray et al. 2021), the contributions of Y1-R may be physiologically significant. As GEP44 acts directly on muscle tissue, while its effects on islets require concomitant Y1-R antagonism, activation of Y1-R expressed in muscle tissue may ultimately prove to be a critical factor in reducing serum glucose levels, alongside any potential effects on insulin secretion and glucagon release.

Our results revealed that Y1-R agonism resulted in insulin-independent glucose uptake into muscle, but only in the presence of elevated glucose levels. (Magnone, Emionite et al. 2020) Although long-term treatment with GEP44 treatment results in profound reductions of serum glucose levels in DIO rats, it stimulates the ISR to a much smaller degree than is observed in response to Ex-4. However, GEP44 co-administered with one of two different Y1-R antagonists resulted in more profound increases in the ISR; these results suggest that Y1-R signaling may activate acute responses of GEP44 mediated by GLP-1R. Both GLP-1R and Y1-R are coupled to the Gα_i_ subunit, in association with reduced levels of cAMP and protein kinase A (PKA) signaling. Given that PKA-mediated signaling is required for glucose-stimulated insulin release, this could be perceived as a contradiction between this mechanism and reports of enhanced insulin secretion under these conditions (Shi, Loh et al. 2015). There are several potential explanations for these observations, including (1) coupling with different G subunits may lead to the activation of protein kinase C (PKC) and ultimately an increase in β-cell mass, especially under conditions of chronic administration, (2) their role in promoting protection against necrotizing or apoptotic β-cell death (Tito, Rudnicki et al. 1993, Sam, Gunner et al. 2012), and (3) activation of phosphatidylinositol 3 kinase y-subunit (PI3Kγ), a kinase identified as a component of the neuropeptide Y signaling cascade which may aid in the correct localization of insulin granules to facilitate insulin secretion (MacDonald 2009); (Goldberg, Taimor et al. 1998).

### Impact of GEP44 on food intake, body weight, and metabolic outcomes in DIO rats compared to responses to Ex-4 and LIRA

Data from the dose escalation experiment conducted in DIO rats indicated that treatment with low doses (0.5 to 10 nmol/kg) of GEP44 and Ex-4 resulted in similar anorectic effects. However, GEP44 has a superior therapeutic window, as rats could be treated with higher doses with no notable adverse effects. As reported in our earlier studies (Milliken, Elfers et al. 2021), GEP44 treatment elicited no indicators of malaise in rats at doses as high as 100 nmol/kg/day. Testing was discontinued at this dose due to extreme reductions in food intake. When comparing the results from the current dose escalation experiment in DIO Wistar rats to those from our earlier experiments performed with lean rats (Milliken, Elfers et al. 2021), and found that GEP44 treatment results in comparable anorectic effects in both models. By contrast, the data presented in this study suggest that Ex-4 treatment results in a more robust anorectic effect in DIO rats compared to their lean counterparts.

Findings from an initial 16-day experiment performed in DIO rats revealed that GEP44 treatment resulted in more profound reductions of food intake and body compared to treatment with equimolar doses (10 and 25 nmol/kg/day) of LIRA, which is a well-established GLP-1RA currently approved for the treatment of obesity. Similar results were observed in a follow-up 27-day treatment study, in which GEP44 continued to elicit more profound reductions in food intake than equimolar doses of LIRA (5, 10, and 25 nmol/kg/day). As part of this longer 27-day treatment study, pair-fed controls were used to determine whether GEP44 or LIRA had any impact on energy expenditure (EE) and the potential to induce weight loss beyond what would be anticipated from reduced food intake alone. Interestingly, mean body weight reductions observed in the two pair-fed groups of DIO rats were consistently lower throughout the experiment compared to those treated with GEP44 and LIRA; however, no significant differences in change of body weight were identified between either of the treated and their respective pair-fed groups. These results are consistent with earlier studies in humans in which LIRA had no impact on EE (Harder, Nielsen et al. 2004); however, current findings do not rule out the possibility of long-term increases in EE associated with drug treatment (Maciel, Beserra et al. 2018).

*Preliminary in vivo studies of GEP12 on food intake and body weight compared to GEP44 in DIO rats* Initial *in vivo* testing with GEP12 (5 and 10 mg/kg/day) resulted in robust reductions in food intake and body weight, albeit somewhat less than responses observed in rats treated with equivalent doses of GEP44. As in prior experiments with GEP44, no indicators of nausea or malaise (e.g., changes in responsiveness, behavior, coat appearance, or facial expressions such as orbital tightening and nose/cheek flattening) were observed with GEP12 dosing.

## CONCLUSION

In summary, the results presented in this manuscript provide pre-clinical validation of a poly-agonistic chimeric peptide targeting GLP-1R, Y1-R and Y2-R. Our findings demonstrate their metabolic stability and selective agonism at Y1-R, Y2-R, and GLP-1R that lead to stimulation of insulin secretion from pancreatic islets and muscle glucose uptake *in vitro* and profound reductions in food intake and body weight in experiments performed *in vivo.* Subsequent work will focus on the use of lipidated analogs of GEP44 and the possibility of one-per-week dosing. This is a promising and innovative route toward the development of unimolecular peptide drugs with superior efficacy than those currently available for the treatment of obesity and T2DM.

## AUTHOR CONTRIBUTIONS

R.P.D. and B.T.M. designed GEP44. R.P.D. and K.S.C. designed GEP12. C.L.R., I.R.S., and R.P.D. developed the study rationale and the experimental designs. K.S.C., C.L.R., I.R.S., and R.P.D. drafted the manuscript, which was reviewed and edited by all authors. All fluorescent probes were designed by K.S.C., B.T.M., and R.P.D. and were synthesized, purified, and characterized by K.S.C., A.M.G., or B.T.M. *In vitro* receptor agonism assays were performed and analyzed by O.G.C. and G.G.H. *Ex vivo* islet and muscle experiments were designed by I.R.S. and conducted by V.K. FISH/RNAScope experiments were performed by S.V.A. and analyzed by S.V.A. and M.R.H. The *in vivo* experiments were performed by C.T.E. and T.S.S. with assistance from K.S.C. and A.M.G. and analyzed by C.T.E. and C.L.R. All authors approved the final version of the manuscript.

### Funding sources, conflicts of interest, and acknowledgments

This work was supported by the United States Department of Defense through a Congressionally Directed Medical Research Program Award (W81XWH1010299) to R.P.D. and C.L.R. and funding from the National Institutes of Health (R01 DK17047, the DRC Cell Function Analysis Core) and the National Science Foundation (STTR 1853066) to I.R.S. G.G.H. received funding via subcontract under National Institutes of Health (R01 DK128443) to R.P.D., M.R.H. and Bart C. De Jonghe. Biochemical analyses were performed at the University of Washington Nutrition and Obesity Research Center (UW NORC) which is supported by grant P30 DK035816 from the National Institute of Diabetes and Digestive and Kidney Diseases. R.P.D. acknowledges the SOURCE program at Syracuse University for funding provided for A.M.G. R.P.D. is a Scientific Advisory Board member of Balchem Corporation, New Hampton, NY, and Xeragenx LLC. (St. Louis, MO); these organizations played no role in the design, execution, or analysis of the results of these studies. R.P.D. and B.T.M. are named authors of a patent pursuant to this work that is owned by Syracuse University. I.R.S. has financial ties to EnTox Sciences, Inc. (Mercer Island, WA), the manufacturer/distributor of the BaroFuse perifusion system used in this study. M.R.H. receives funding from Zealand Pharma, Novo Nordisk, Eli Lilly & Co., and Boehringer Ingelheim; these funds were not used to support these studies. M.R.H. and R.P.D. are co-founders and co-owners of Cantius Therapeutics (Lansdale, PA), which also played no role in these studies.

## STAR*METHODS

Detailed methods are provided in the online version of this paper and include the following:

## Key Resources Table

| REAGENT or RESOURCE | SOURCE | IDENTIFIER |
| --- | --- | --- |
| **Antibodies** |  |  |
| Anti-anti-Cy5 Antibody (B-2) Alexa Fluor 647 | Santa Cruz Biotechnology | sc-166896 AF647 |
| **Bacterial and virus strains** |  |  |
| H188 Adenovirus | Lab of Prof. Kees Jalink, Division of Cell Biology, the Netherlands Cancer Institute, Amsterdam, the Netherlands. | N/A |
| **Biological samples** |  |  |
| Rat Islets | Harvested from Sprague-Dawley rats (n=23; ~250g; Envigo/Harlan) | N/A |
| Human Islets | Human islets were provided by the NIH-funded Integrated Islet Distribution Program | N/A |
| Rat brain tissue | Extracted from adult male Sprague-Dawley rats (n=23; ~400g; Charles River Laboratories) | N/A |
| **Chemicals, peptides, and recombinant proteins** |  |  |
| Exendin-4 | In-house (Syracuse University) | N/A |
| Liraglutide | Selleck Chemicals (Houston, TX) | S8256 |
| GEP44 | GenScript (Piscataway, NJ) | N/A |
| GEP12 | In-house (Syracuse University) | N/A |
| PYY(3-36) | In-house (Syracuse University) | N/A |
| PYY(1-36) | In-house (Syracuse University) | N/A |
| PD160170 | Tocris | 2200 |
| BIBO3304 | Tocris | 2412 |
| <sup>3</sup>H-2-deoxyglucose | Perkin Elmer | NET328A250UC |
| Wortmannin | Sigma Aldrich | W1628 |
| Y1-R | antibodies-online Inc. | ABIN4888949 |
| Glucagon-Like Peptide 1 Receptor (GLP-1R) protein (His tag) | antibodies-online Inc. | ABIN3080888 |
| Neuropeptide Y Receptor Y1 (NPY1R) protein (His tag) | antibodies-online Inc. | ABIN7086253 |
| ProTide Rink amide resin | CEM Corporation | R002 |
| Triisopropylsilane | Sigma-Aldrich | 233781 |
| Trifluoroacetic acid | Sigma-Aldrich | 8.08260.2501 |
| *N,N'*-Diisopropylcarbodiimide | Sigma-Aldrich | D125407 |
| Oxyma Pure | CEM Corporation | S001 |
| Piperidine | Sigma-Aldrich | 8.22299.0500 |
| *N,N'*-Dimethylformamide | VWR (Radnor, PA) | BDH83634.400 |
| α-cyano-4-hydroxycinnamic acid | Acros Organics | 163440050 |
| Acetonitrile HiSolv | VWR | BDH83639.400 |
| Diethyl ether | VWR | BDH67003.400 |
| Dulbecco's Modified Eagle Medium | Sigma-Aldrich | D6429 |
| Penicillin-streptomycin | ThermoFisher Scientific | 15140122 |
| Bovine serum albumin | Sigma-Aldrich | A3059 |
| Fetal bovine serum | Sigma-Aldrich | 12303C |
| Liberase | Roche Molecular Biochemicals (Indianapolis, IN) | 05339880001 |
| Euthasol, 390 mg/ml sodium pentobarbital | Virbac | RXEUTHASOL |
| Insulin | MilliporeSigma | 91077C |
| Catheter lock solution (500 USP units/ml heparin in 50% glycerol) | Instech Labs (Plymouth Meeting, PA) | USP-HGS-500-10-VBP-5 |
| DAPI-containing mounting media | VECTASHIELD Antifade Mounting Medium | H1200 |
| Lactate | MilliporeSigma | L9795 |
| Standard Extracellular Saline (SES) Solution | In-house (Upstate Medical University) | N/A |
| Normal donkey serum | Sigma Aldrich | 566460 |
| 5-thio-D-glucose | Santa Cruz Biotechnology | sc-221044A |
| Meloxicam | Midwest Veterinary Supply | N/A |
| isoflurane | Butler Schein |  |
| sulfo-Cyanine5 DBCO | Lumiprobe | 433F0 |
| Bay-K8644 | Sigma-Aldrich | B112 |
| Critical commercial assays and kits |  |  |
| hY2-R Binding Assay | EuroscreenFast (Gosselies, Belgium) | FAST-0321B |
| hGLP-1R Agonist-Based Internalization | Eurofins Discovery (Fremont, CA) | 86-0010P-2029AG |
| hY2-R Agonist-Based Internalization | Eurofins Discovery (Fremont, CA) | 86-0010P-2037AG |
| hGLP-1R Agonist-Based Arrestin Recruitment | Eurofins Discovery (Fremont, CA) | 86-0001P-2166AG |
| Leptin, IL-1β, IL-6, and TNF-α Magnetic Bead Panel Assay | MilliporeSigma | RECYTMAG-65K |
| Insulin ELISA | MilliporeSigma | EZRMI-13K |
| Adiponectin ELISA | MilliporeSigma | EZRADP-62K |
| cAMP Kit | ThermoFisher | 4412182 |
| Amplex Red Glucose/Glucose Oxidase Assay Kit | ThermoFisher | A22189 |
| Lactate Oxidase from *Aerococcus viridans* | MilliporeSigma | L9795 |
| Beuthanasia-D | Schering-Plough Animal Health Corp., Union, NJ | N/A |
| Experimental models: Cell lines |  |  |
| HEK293 cells stably transfected to express hGLP-1R | In-house | N/A |
| HEK293 cells | American Type Culture Collection (Manassas, VA) | N/A |
| HEK293 C24 | American Type Culture Collection (Manassas, VA) | N/A |
| Experimental models: Organisms/strains |  |  |
| Adult male Wistar rats | Charles River Laboratories (Wilmington, MA) | 003 |
| Adult male Sprague Dawley rats | Envigo Harlan (Indianapolis, IN) | 002 |
| Adult male DIO Wistar rats | Charles River Laboratories<br>(Wilmington, MA) | 003 |
| Software and algorithms |  |  |
| GraphPad PRISM | GraphPad Software | N/A |
| Stata/SE 14.2 | STATA Corp LLC | N/A |
| ProData Viewer Software | JASCO J-715 spectropolarimeter | N/A |
| FlexStation 3 microplate reader | Molecular Devices | N/A |
| Imaris 8.1.2 software | Bitplane | N/A |
| HPEPDOCK Server | Huang Lab (Huazhong University<br>of Science and Technology,<br>Wuhan, China) | N/A |
| Other |  |  |
| High-Fat diet | Research Diets, Inc. (New<br>Brunswick, NJ) | D12492 |
| BioDAQ cages | Research Diets, Inc. (New<br>Brunswick, NJ) | E2 Electronic |
| DietMax Food Monitoring System | OmniTech Electronics, Inc.<br>(Columbus, OH) |  |
| Normal chow | LabDiet (St. Louis, MO) | PicoLab Rodent 5053 |
| Kaolin (powdered) | Sigma Aldrich | K1512 |
| OneTouch Ultra Mini Glucometer | Lifescan (Malvern, PA) |  |
| OneTouch Ultra Glucose test strips | Lifescan (Malvern, PA) | N/A |
| Microvette® 100 K<sub>3</sub> EDTA | Sarstedt | 20.1278.100 |
| Superfrost Plus slides | Fisher Scientific |  |
| RNAscope® Multiplex Fluorescent Reagent Kit v2 | ACDBio | 323100 |
| RNAscope probe Rn-NPY1-R-C1 | ACDBio | 414471 |
| RNAscope probe Rn-GLP-1R-C2 | ACDBio | 315221-C2 |
| RNAscope probe Rn-NPY2-R-C1 | ACDBio | 414481 |
| BZ-X800 microscope | Keyence |  |
| Liquid scintillation counter | Beckman | Model LS6500 |
| 26-gauge cannula | Plastics One |  |
| Spectrophotometer, plate reader, Synergy 4 | BioTek (Winooski, VT) | Model S4MLFPTA |
| Wizard 2, 5-channel gamma counter | Perkin Elmer | Model 2470-0050 |
| BaroFuse Multi-Channel perfusion system | EnTox Sciences | Model 001-08 |
| pcDNA3.1-hY2 receptor plasmid DNA | cDNA Resource Center | NPYR20TN00 |
| pcDNA3.1-hY1 receptor plasmid DNA | cDNA Resource Center | NPYR10TN00 |
| OpenSPR | Nicoya (Kitchener, ON, Canada) | N/A |
| Nitrilotriacetic acid (NTA) sensor chip | Nicoya Store (Kitchener, ON,<br>Canada) | SEN-AU-100-10-NTA |
| Zorbax C18 column (5µm, 9.4 x 250 mm) | Agilent | 880995-202 |

## Resource Availability

### Lead Contact

Further information and requests for resources and reagents should be directed to and will be fulfilled by the lead contact Professor Robert P. Doyle.

### Materials Availability

This study generated new unique reagents. R.P.D is the named author of a patent pursuant to this work that is owned by Syracuse University and will supply the reagent under MTA upon reasonable request.

### Data and Code Availability

The published article includes all data generated or analyzed during this study. No code was used or generated in this study.

## METHOD DETAILS

### Peptide Syntheses and Purification

Solid-phase peptide synthesis was performed on ProTide Rink amide resin using a microwave-assisted CEM Liberty Blue peptide synthesizer (Matthews, NC, USA). Fmoc-protected amino acids were coupled to the resin using Oxyma Pure (0.25 M) and N, N’-diisopropyl carbodiimide (0.125 M) as the activator and activator base, respectively. Fmoc was removed between couplings with 20% piperidine. Global deprotection and cleavage of the peptides from the solid-support resin were achieved using a CEM Razor instrument via a 40-minute incubation at 40°C in a mixture of 95% trifluoroacetic acid, 2.5% triisopropylsilane, and 2.5% water. Peptides were precipitated with cold (4°C) diethyl ether and purified on an Agilent 1200 series High-Performance Liquid Chromatography (HPLC) instrument (10–75% HPLC-grade acetonitrile for 20 minutes at a 2 mL/min flow rate over an Agilent Zorbax C18 column (5 µm, 9.4 x 250 mm) tracked at 220, 254, and 280 nm. Peptides were purified to >95%. Peptide binding was assayed by EuroscreenFast (Gosselies, Belgium) or in-house with a Nicoya Open Surface Plasmon Resonance (SPR) instrument. Internalization and ý-arrestin recruitment assays were performed by Eurofins Discovery (Fremont, CA, USA).

### Competitive Binding Assays at GLP-1R

GEP44 and Ex-4 binding to the human GLP-1R was measured using a TagLite fluorescent competitive binding assay in CHO-K1 cells. GLP-1*_red_* was used as the agonist tracer and Ex-4 as the reference competitor. IC_50_ values were measured in duplicate in independent runs at eight concentrations per run. GEP12 binding to the human GLP-1R was measured in-house by SPR using His-tagged GLP-1R bound to a nitrilotriacetic acid (NTA) sensor. The GEP12 dose-response (0.1 nM – 100 nM) binding assay was performed in a duplicate.

### Competitive Binding Assays at Y2-R

Peptide binding to the human Y2-R was measured in a dose-responsive manner (1 pM –1 µM) using a radioligand competitive binding assay in CHO-K1 cells. Peptide binding was assayed in duplicate independent runs with eight concentrations per run. The peptide PYY_3-36_ was used as a positive control.

### Competitive Binding Assays at Y1-R

Peptide binding to human Y1-R was performed in-house by SPR. The dose response to GEP44 (4 pM - 19 µM) was evaluated using PYY_1-36_ as positive and PYY_3-36_ as negative controls.

### Internalization of GLP-1R

Human GLP-1R (G_s_-coupled) cell-based agonist-activated internalization assays were performed by Eurofins Discovery. Dose-response assays (30 pM - 1 µM) were performed in duplicate with Ex-4 as a positive control.

### Internalization of Y2-R

A human Y2-R (G_s_-coupled) dose-response (5.51 pM - 551 nM) cell-based agonist-activated internalization assay was performed in duplicate with PYY_3-36_ as a positive control.

### β-Arrestin Recruitment at GLP-1R

Human GLP-1R cell-based arrestin assays were performed by Eurofins Discovery (Fremont, CA, USA; assay #86-0001P-2166AG) as described by the company. Dose-response assays (5.51 pM – 551 nM) were performed in duplicate with Ex-4 as a positive control.

### *In vitro* Receptor Agonism at Y1-R, Y2-R, and GLP-1R

H188 virally transduced HEK293 cells stably expressing human GLP-1R were obtained from Novo Nordisk A/S for use in FRET assays. HEK293 C24 cells stably expressing the H188 FRET reporter were obtained by G418 selection and grown in monolayers to ∼70% confluency in 100 cm^2^ tissue culture dishes and were then transfected with plasmids (11 μg/dish) encoding human GLP-1R, human Y2-R, or human Y1-R. Transfected cells were then incubated for 48 h in fresh culture media. For real-time FRET kinetic assays, cells were harvested, resuspended in 21 mL of SES buffer, and plated at 196 μL per well. Plated cells were pretreated with 4 μL of agonist, or antagonist (Ex_9-39_ (GLP-1R antagonist) or BIIE0246 (Y2-R antagonist)), at a given target concentration and incubated for 20 min before performing the assay. Y1-R and Y2-R agonism to stimulate G_i_ proteins and to inhibit adenylyl cyclase was monitored by detecting the ability of PYY peptides, GEP44, or GEP12 to counteract the ability of Adenosine (acting through endogenous A2B receptor and Gs proteins) to increase levels of cAMP. For these assays, increased levels of cAMP were measured as an increase of the 485/535 nm FRET ratio serving as a readout for binding of cAMP to the H188 biosensor that is based on the exchange protein activated by cAMP (Chepurny, Bonaccorso et al. 2018).

#### Rat islet isolation and culture

Islets were harvested from Sprague-Dawley rats (∼250 g) that were anesthetized by an intraperitoneal injection of pentobarbital sodium (150 mg/kg). All procedures were approved by the University of Washington Institutional Animal Care and Use Committee (IACUC Protocol 4091-01). Islets were prepared and purified as described (Rountree, Neal et al. 2014). Briefly, islets were prepared by injecting collagenase (10 mL of Liberase at 0.23 mg/mL) into the pancreatic duct followed by surgical removal of the pancreas. The isolated pancreata were placed into 15 mL conical tubes containing 10 mL of 0.23 mg/mL Liberase and incubated at 37°C for 30 min. The digests were then filtered and rinsed with Hank’s buffered salt solution (HBSS). Islets were purified using an Optiprep gradient (Nycomed, Oslo, Norway) as previously described (Brandhorst, Brandhorst et al. 1999) and cultured for 18 h in a 37°C in a 5% CO_2_ incubator in Roswell Park Memorial Institute (RPMI) medium supplemented with 10% heat-inactivated fetal bovine serum before use in experiments.

#### Static measurements to determine rates of insulin secretion, glucagon secretion, and cAMP release

Rates of insulin, glucagon, and cAMP release were determined statically under multiple conditions as previously described (Jung, Reed et al. 2009). Briefly, islets were handpicked, transferred to a petri dish containing 11 mL of Krebs-Ringer bicarbonate (KRB) buffer supplemented with 0.1% bovine serum albumin (BSA) and 3 mM glucose, and incubated at 37°C and 5% CO_2_ for 60 min. Islets were then selected and transferred into wells of 96-well plates containing 0.2 mL of KRB with 20 mM glucose and various test compounds as indicated and incubated for an additional 60 min. Subsequently, the supernatants were assayed for insulin, glucagon, and cAMP. These values were used to calculate the secretion rate as the concentration in the assay (ng/ml for insulin and glucagon, and pmol/mL for cAMP) times the volume of KRB in each well during the assay (0.2 mL) divided by the assay time (60 minutes). Data was then normalized by dividing by the secretion rate in the presence of test compounds over that obtained at 20 mM glucose alone.

#### Preparation of test compounds for *in vitro* experiment

Adding test compounds to solutions for either static or perifusion protocols involved making up a stock solution and then adding a small volume to the wells (for static) or inflow buffer (for perifusion). Stock solutions of test compounds that are water soluble were made up at 20 times the final assay concentration in buffer (including glucose, GEP44, GEP12, exendin-4, PYY_1-36_ and insulin). For test compounds that were insoluble in water (Y1-R antagonists) stocks were made up at 1000 times the assay concentration in DMSO, so that final concentration of DMSO in the assay was 0.1%.

#### Perifusion measurements to determine rates of insulin secretion

Insulin production was evaluated using a commercially available perifusion system (BaroFuse; EnTox Sciences, Mercer Island, WA). Ten isolated rat islets were placed into each of eight channels that were operating at a flow rate of 50 μL/min of KRB (continuously equilibrated with 21% O_2_ and 5% CO_2_ and balance of N_2_) containing 0.1% BSA and 3 mM glucose for 90 minutes. Subsequently, varying amounts of glucose and test compounds were injected into the inflow of the flow system as indicated and outflow fractions were collected every 10 minutes and assayed for insulin as described in the section to follow. The insulin secretion rate was calculated as insulin times the flow rate of KRB divided by the number of islet x 100 yielding the ng/min/100 islets. Data was graphed after normalizing each insulin time course by dividing by the secretion rate at each time point by the rate obtained in the presence of 20 mM glucose prior to the addition of test compounds.

#### Assays performed on supernatants collected from static incubation and outflow fractions of perifusion experiments

Assays using commercially available kits were performed according to the manufacturers’ instructions.

#### Insulin measured by radioimmunoassay (RIA)

Briefly, specific anti-insulin antiserum was incubated with the sample together with defined amounts of ^125^I-labeled insulin. Antibody-bound tracer was separated from the unbound tracer by precipitation in solution provided in the kit containing 3% PEG and 0.05% Triton X-100 in 0.05M Phosphosaline with 0.025M EDTA and 0.08% sodium azide. The ^125^I remaining in the tube was assessed quantitatively on a five-channel gamma counter. The amount of ^125^I detected in the supernatant is inversely proportional to the amount of insulin in the original sample.

#### Glucagon measured by immunoassay

Briefly, samples were incubated with two manufacturer-supplied anti-glucagon monoclonal antibodies that were covalently linked to either SmBiT or LgBiT. After the detection substrate was added, the resulting luminescence was measured using a spectrophotometer. The luminescent signal detected is directly proportional to the amount of glucagon present in the sample.

#### Quantitative evaluation of cAMP levels by ELISA

Samples and diluted cAMP-alkaline phosphatase (AP) reagent were added to wells of a pre-coated assay plate (Thermofisher; Waltham, MA) and mixed by repetitive pipetting. After a one-hour incubation, the solutions were removed from wells which were then washed six times with wash buffer. A substrate/enhancer solution was then added, and the plates were incubated for 30 min. The luminescent signal was measured using a spectrophotometer at room temperature protected from light.

#### Glucose uptake in muscle

^3^H-2-deoxyglucose (DG) uptake into muscle tissue was evaluated as previously described (Sweet, Cook et al. 2004), with the exception that the bound radiolabeled compound was separated from free radiolabel by washing the tissue multiple times in radiolabel-free medium. Sprague-Dawley rats (∼250 g) were anesthetized by intraperitoneal injection of Beuthanasia-D (38 mg pentobarbital sodium and 6 mg phenytoin sodium/230 g rat) (Schering-Plough Animal Health Corp., Union, NJ). While the rats remained under anesthesia, strips of quadriceps muscle were collected and transferred to a Petri dish containing HBSS with 0.1% BSA. While still under anesthesia, animals were then euthanized by cutting the diaphragm. The muscle strip was cut into smaller pieces (∼2 mg each) using a scalpel. Three pieces were then transferred into polystyrene 12 x 75 test tubes containing 190 mL of KRB (with 5 mM bicarbonate) solution and compounds as described in each experiment. Each condition was evaluated in triplicate. The tubes were placed in racks that were partially submerged in a shaking water bath maintained at 37°C. At precisely the times indicated, 10 μL of the radioactive dose (typically 0.25 μCi) was spiked into each tube to bring the final volume to 200 mL. The tubes were capped and shaken in the water bath at 120 rpm for precisely 45 minutes. Free radiolabel was removed by washing the muscle fragments three times with 5 mL cold (4°C) KRB solution. After the third wash, 100 μL of KRB was added to each tube and the muscle and solution were then transferred to a microcentrifuge tube. The muscle pieces were then fragmented further by sonication (Branson) at maximum power and 50% duty cycle for 20 seconds. The contents of the microfuge tube were then transferred to a 7 mL scintillation vial. A liquid scintillation cocktail (5 mL, Ecolume) was added, the samples were shaken, and radioactivity was evaluated using a liquid scintillation counter

### Lactate production by perifused muscle tissue

Muscle fragments (6 x 2 mg each) were placed into each of six channels of a commercially available perifusion system (BaroFuse) operating at a flow rate of 30 μL/min of KRB with 0.1% BSA and varying amounts of glucose. Outflow fractions were collected every 10 minutes and measured using a glucose/glucose oxidase kit in which lactate oxidase was used to replace glucose oxidase. Manufacturer-supplied solutions of horseradish peroxidase, Amplex Red, and lactate oxidase were added to samples in wells of a 96-well microplate which was then incubated at room temperature for 30 minutes. Fluorescence was measured with a spectrophotometer.

### Fluorescent *in situ* hybridization (FISH) and immunohistochemical (IHC) visualization of fluorescent GEP44 (f-Cy5-GEP44) and its localization AP/NTS neurons that express GLP-1R and Y1-R or Y2-R

Male Sprague-Dawley rats were briefly anesthetized with isoflurane (5% induction followed by 2–3% maintenance) to facilitate implantation of an in-dwelling cannula (26 gauge) directed at the 4^th^ cerebroventricular region (coordinates: on the midline, 2.5 mm anterior to the occipital suture and 5.2 mm ventral to the skull.) Postoperative analgesia (2 mg/kg meloxicam) was administered subcutaneously for two days, and the rats were allowed to recover for one week. Proper placement and cannula patency were verified via 5-thio-D-glucose (210 μg)-induced hyperglycemia, as previously described (Mietlicki-Baase, Liberini et al. 2018). Rats with appropriate cannula placement and patency were included in the experiments to follow. Rats (n=2) received an ICVI of f-Cy5-GEP44 (1 μg) dissolved in 1 μL 0.9% normal saline solution. Additional rats (n=2) received an intraperitoneal injection of f-Cy5-GEP44 (15.5 μg/kg). One hour later, rats were anesthetized with ketamine (90 mg/kg), xylazine (2.8 mg/kg), and acepromazine (0.72 mg/kg) and transcardially perfused with 0.1 M phosphate-buffered saline (PBS) followed by 4% paraformaldehyde (PFA) in PBS. Brains were collected and stored in 4% PFA for 24 hours after which they were transferred to 20% sucrose for cryoprotection at 4°C. Serial coronal sections (16 μm thickness) of each brain were prepared using a cryostat, mounted onto Superfrost Plus slides, and stored at −80 °C. One series that included the rostral-caudal extent of the NTS was used to detect RNA levels of Y1-R (RNAscope Probe Rn-NPY1-R-C1), GLP-1R (RNAscope Probe Rn-GLP-1R-C2), and f-Cy5-GEP44; another series was used separately to detect RNA levels of Y2-R (RNAscope Probe Rn-Y2-R-C1), GLP-1R, and f-Cy5-GEP44. Fluorescent *in situ* hybridization (FISH) was performed as per the manufacturer’s instructions for the RNAscope® Multiplex Fluorescent Reagent Kit v2. Sections were washed three times for 5 min each in 0.1 M PBS, followed by incubation in 50% ethanol for 30 min. Sections were then rinsed again (three times x 5 min each with 0.1 M PBS). Sections were then washed in freshly prepared 0.1% sodium borohydride for 20 min and washed again in 0.1 M PBS (three times, 5 min each). To restore Cy5-GEP44 fluorescence, sections were then incubated overnight with an Alexa Fluor 647-linked mouse anti-Cy5 antibody (1:100). After washing (three times x 5 min each with 0.1 M PBS, the tissues were immersed in DAPI-containing mounting media, cover-slipped, and stored at 4 °C. Images were acquired 24 – 48 hrs later, on a Keyence BZ-X800 microscope using negative control sections to adjust for background fluorescence. Images were taken using filter cubes for DAPI, GFP, Cy3, and Cy5 at 20x and 40x magnification. Image z-stacks were collected with a step-size of 1 μm and rendered as three-dimensional rotational animations using Imaris 8.1.2 software (Supplementary videos 1 and 2).

### Animal experiments

All procedures performed in rats were approved by the Institutional Animal Care and Use Committee at the Seattle Children’s Research Institute (IACUC00064) and were in accordance with the National Institutes of Health (NIH) Guide for Care and Use of Laboratory Animals. This facility is approved by the Association of the Assessment and Accreditation of Laboratory Animal Care International (AAALAC). Wistar rats purchased from Charles River Laboratories (Wilmington, MA) were used in this study. The rats were provided with *ad libitum* access to food and water and were kept on a 12 h light/12 h dark cycle. For all other experiments, male rats (51–75g; approximately 4 weeks of age) were fed a diet with 60% of the calories provided by fat (D12492; Research Diets, Inc; New Brunswick, NJ; 5.21 kcal/g) for approximately five months before the start of the study generate diet-induced obesity (DIO). Animals were individually housed in a temperature (22 ± 1 °C) and humidity-controlled (57 ± 4%) room. All body weight measurements were taken just before the start of the dark cycle.

### Preparation and administration of drugs

GEP44 stock solutions were prepared using sterile ultra-pure H_2_O and gently agitated at 4°C for >24 h and then aliquoted and stored at -20°C. Ex-4 was produced by the Doyle lab and dissolved in 1 mL of sterile ultra-pure H_2_O. LIRA was dissolved in 1 mL of sterile ultra-pure H_2_O. The stock solutions were aliquoted and stored at -20°C. Stock solutions for both Ex-4 and LIRA were diluted to working solutions in 0.9% normal saline and mixed gently before use. GEP12 (aliquots up to 5 mg) was dissolved in 100 μL DMSO and mixed at 1000 rpm for three min to create a stock solution. Stock solutions were diluted to working solutions in 0.9% normal saline, resulting in a final DMSO concentration of <2%. The unused portion of the stock solution was stored at -20°C for up to 7 days. Working solutions were stored at 4°C for up to two days and mixed gently before use. Normal saline solution was used as the vehicle control in all studies. Working solutions of GEP44, GEP12, Ex-4, LIRA, or vehicle control were administered by subcutaneous injection once daily at the start of the dark cycle using a 1 cc 29G insulin syringe. The concentration of each drug solution was adjusted to maintain the dosing volume at 0.5 mL/kg.

### Dose escalation study in diet-induced obese male rats

Male Wistar rats (Ex-4 group, n=4; GEP44 group, n=8) were provided with a high-fat (HF) diet (60% cal from fat, 5.21 kcal/g) for 20 weeks before the start of the study. Rats were singly housed in BioDAQ cages and allowed to acclimate for at least 10 days before the start of this study. The average baseline weight of the rats in this study was 685 ± 38 g. Baseline measurements of body weight and food intake were taken for three days to balance the groups. The study design included sequential rounds of a three-day vehicle-treated baseline phase, a three-day treatment phase, and a two-to-three-day washout phase. Dosing began at 0.5 nmol/kg/day and was increased in approximately one-third-log increments (10^n/3^) until the MTD was established. The doses tested included 0.5, 1, 2, 5, 10, 20, 50, and 100 nmol/kg/day administered subcutaneously just before the start of the dark cycle. The 100 nmol/kg dose of GEP44 was tested for one day only. Body weight was assessed daily just before the start of the dark cycle. Food and water were available *ad libitum* and consumption was monitored continuously.

A preliminary GEP12 dosing test was performed in male DIO Wistar rats (n=8). These rats were fed the HF diet for 40 weeks before the start of the study and weighed an average of 862 ± 82 g at baseline. The study design included two rounds of a three-day vehicle-treated baseline phase, a three-day treatment phase, and a two-to-three-day washout phase. Doses of 5 to 10 nmol/kg/day were administered via subcutaneous injection just before the start of the dark cycle. Body weight was assessed daily just before the start of the dark cycle. Food and water were available *ad libitum* and consumption was continuously monitored using a BioDAQ system. In this experiment, cages were modified to facilitate the use of the DietMax food monitor system for continuous recording of powdered kaolin consumption. Animals were allowed to acclimate for at least 10 days before the start of the study.

### Long-term drug intervention

Male Wistar rats (n=4) were fed a 60% HF diet for 20 weeks before the start of the study. Rats were then housed singly in BioDAQ cages and allowed to acclimate to their new environment for 10 days. Baseline measures of body weight and food intake were collected for one week to balance the groups and create feeding pairs. Two independent experiments were performed. In the first experiment, three cohorts of eight animals each received daily injections of vehicle or increasing doses of either GEP44 or LIRA starting at 10 nmol/kg for 9 days and followed by 25 nmol/kg for 7 days. Rats averaged 802 ± 86 g at baseline with equivalent variances between the groups. In the second experiment, five cohorts of eight animals each received daily injections of GEP44 alone, vehicle pair-fed with GEP44, LIRA alone, vehicle pair-fed to LIRA, and vehicle alone. Rats averaged 661 ± 68 g at baseline with equivalent variances between the groups. GEP44 was administered as follows: 5 nmol/kg/day for 4 days, 10 nmol/kg/day for 4 days, 25 nmol/kg/day for 12 days, and 50 nmol/kg/day for 8 days. LIRA was administered as follows: 5 nmol/kg/day for 4 days, 10 nmol/kg/day for 4 days, 25 nmol/kg/day for 4 days, and 50 nmol/kg/day for 16 days. Food intake was monitored continuously throughout the experiment. Body weights were measured daily immediately before the start of the dark cycle.

Pre- and post-treatment fasting plasma samples were obtained from blood collected via tail nick using a microvette to assess insulin levels. Blood glucose concentrations were obtained at the same time using a handheld glucometer. Blood was collected by cardiac puncture at the time of euthanasia which was two hours after the final injection. Commercially available enzyme-linked immunosorbent assays (ELISAs) were used to perform quantitative assessments of both insulin and adiponectin. A commercially available magnetic bead panel was used to perform quantitative assessments of leptin, IL-1β, IL-6, and TNF-α. Serum samples were diluted 1:500 for the adiponectin ELISA using the sample diluent provided with the kit. All ELISAs were performed following the manufacturers’ recommendations. Serum levels of glucose, cholesterol (total, high-density lipoprotein [HDL], and calculated low-density lipoprotein [LDL]), triglycerides, alanine transaminase (ALT), and aspartate transaminase (AST) were determined using a Modular P chemistry analyzer (Roche Diagnostics, Germany) by the University of Washington NORC Core, Seattle, WA.

## QUANTIFICATION AND STATISTICAL ANALYSIS

All data were expressed as mean ± SD unless otherwise noted. For behavioral studies, data were analyzed by ANCOVA, or repeated-measures one-way or two-way ANOVA followed by Tukey’s or Bonferroni’s post hoc test as appropriate. For all statistical tests, a *p*-value <0.05 was considered significant. All data were analyzed using Prism GraphPad 9 or Stata/SE 14.2.

